# Oncogenic KEAP1 mutations activate TRAF2-NFκB signaling to prevent apoptosis in lung cancer cells

**DOI:** 10.1101/2023.10.10.561664

**Authors:** Ashik Jawahar Deen, Simone Adinolfi, Jouni Härkönen, Tommi Patinen, Xiaonan Liu, Tuomo Laitinen, Piia Takabe, Kirsi Kainulainen, Sanna Pasonen-Seppänen, Lisa M Gawriyski, Uma Thanigai Arasu, Ilakya Selvarajan, Petri Mäkinen, Hanna Laitinen, Emilia Kansanen, Minna U Kaikkonen, Antti Poso, Markku Varjosalo, Anna-Liisa Levonen

## Abstract

The Kelch-like ECH-associated protein 1 (KEAP1) – Nuclear factor erythroid 2-related factor 2 (NRF2) pathway is the major transcriptional stress response system in cells against oxidative and electrophilic stress. NRF2 is frequently constitutively active in many cancers, rendering the cells resistant to chemo- and radiotherapy. Loss-of-function (LOF) mutations in the repressor protein KEAP1 are common in non-small cell lung cancer, particularly adenocarcinoma. While the mutations can occur throughout the gene, they are enriched in certain areas, indicating that these may have unique functional importance. In this study, we show that in the GSEA analysis of TCGA lung adenocarcinoma RNA-seq data, the KEAP1 mutations in R320 and R470 were associated with enhanced Tumor Necrosis Factor alpha (TNFα) *–* Nuclear Factor kappa subunit B (NFκB) signaling as well as MYC and MTORC1 pathways. To address the functional role of these hotspot mutations, affinity purification and mass spectrometry (AP-MS) analysis of wild type (wt) KEAP1 and the mutants was employed to interrogate differences in the protein interactome. We identified TNF receptor associated factor 2 (TRAF2) as a putative protein interaction partner. Both mutant KEAP1 forms showed increased interaction with TRAF2 and other anti-apoptotic proteins, suggesting that apoptosis signalling could be affected by the protein interactions. A549 lung adenocarcinoma cells overexpressing mutant KEAP1 showed high TRAF2-mediated NFκB activity and increased protection against apoptosis, XIAP being one of the key proteins involved in anti-apoptotic signalling. To conclude, KEAP1 R320Q and R470C and its interaction with TRAF2 leads to activation of NFκB pathway, thereby protecting against apoptosis.

## Introduction

The KEAP1 – NRF2 pathway is a major transcriptional stress response system in cells against oxidative, chemical and other environmental insults (Cuadrado *et al*, 2019; Pillai *et al*, 2022). KEAP1 acts as a repressor for NRF2 under basal conditions, and NRF2 is ubiquitinated for proteasomal degradation by the KEAP1-Cullin3-E3 ligase complex (Deen *et al*, 2018; Patinen *et al*, 2019; Jaramillo & Zhang, 2013). In many cancers, non-small cell lung cancer (NSCLC) in particular, NRF2 is constitutively active promoting growth and rendering cells resistant to chemo- and radiotherapy (Okazaki *et al*, 2022; Silva *et al*, 2019; Matsuoka *et al*, 2022). This can occur via inactivation of KEAP1 by several mechanisms such as loss-of-function (LOF) mutations, p62 mediated degradation by autophagy, and post-translational modifications resulting in constitutive activation of NRF2 (Leinonen *et al*, 2015a). Also, NRF2 can have activating mutations/amplifications especially in squamous cell cancers, supporting NRF2 activity (Sanchez-Vega *et al*, 2018; Leinonen *et al*, 2015b; Kansanen *et al*, 2013; Adinolfi *et al*, 2023; Härkönen *et al*, 2023). In NSCLC, KEAP1 and NRF2 are among the most significantly mutated genes, illustrating the importance of the KEAP1-NRF2 pathway in its pathogenesis (Campbell *et al*, 2016). Most of the mutations in both genes are point mutations resulting in a change in a single amino acid residue (Romero *et al*, 2020). Although KEAP1 LOF mutations can occur anywhere, certain sites are mutation hotspots, motivating research on their functional impact (Hast *et al*, 2013, 2014). There are two KEAP1 mutations in particular, R320Q and R470C, which are interesting due to their ability to bind more avidly to NRF2 (Hast *et al*, 2014). Though these mutations do not affect the ubiquitination of NRF2, it is assumed that this increased binding could lead to competitive inhibition thereby slowing down the rate of NRF2 degradation (Hast *et al*, 2014).

NRF2 signaling is regulated by proteins that disrupt the KEAP1-NRF2 binding interface. A number of KEAP1 interacting proteins such as P62, DPP3 and iASPP have been shown to be involved in the inhibition of KEAP1-mediated ubiquitination and proteasomal degradation of NRF2, resulting in NRF2 stabilization and transactivation of target genes (Hast *et al*, 2013; Ge *et al*, 2017). In addition, tumor suppressor proteins WTX, CDKN1A and PALB2 can competitively bind to KEAP1 in a similar manner (Camp *et al*, 2012; Chen *et al*, 2009; Tamberg *et al*, 2018). To explore KEAP1 interacting proteome in an unbiased manner, affinity purification and mass spectrometry (AP-MS) has been used (Hast *et al*, 2013, 2014). Identified proteins include well known KEAP1 interacting proteins P62, PGAM5 and DPP3 but has also revealed novel interacting partners of functional importance such as MCM3 (Yang *et al*, 2022), illustrating the power of this approach.

In this study, we have evaluated hot spot mutations of KEAP1 from the TCGA lung adenocarcinoma cohort, correlating differential expression of genes with the respective mutations. In gene set enrichment analysis (GSEA), KEAP1 mutations that showed increased binding ability to NRF2 were enriched in the TNFα-NFκB signaling along with cell cycle, mTORC1 and MYC pathways. We also employed AP-MS to interrogate KEAP1 interacting proteins and identified TRAF2 as a prominent interacting partner of wt KEAP1 and its mutant forms R320Q and R470C, with TRAF2 binding more avidly to the mutant forms. Given that TRAF2 is the key signaling protein modulating NFκB activity and apoptotic signaling, we also explored how these pathways are altered by mutant KEAP1.

## Results

### 1. TCGA lung adenocarcinoma samples with KEAP1 driver mutations form three unique clusters and show differential gene expression in oncogenic signaling pathways

In cBioportal database, we analyzed KEAP1 mutations in NSCLC from a total of 6887 patients in 20 different studies, including TCGA pan-cancer atlas. Genetic alterations such as point mutations, truncation and copy number alterations were identified in 13% of patients. Somatic mutations contributed to 10.9% (751/6887 patients) of the altered cases, out of which 40.9% (256/626 patients) and 19.3% (121/626 patients), respectively, constitute driver and missense driver mutations (cBioportal.org). R320, R470 and G333 were among the most frequently mutated sites in KEAP1 and were recognized as putative missense driver mutations in NSCLC (Fig. 1A). To characterize the KEAP1 driver mutations and study their effect on gene expression, we next investigated the RNA sequencing data from the available TCGA lung adenocarcinoma cohort. Principal Component Analysis (PCA) of the RNA sequencing data revealed that KEAP1 mutants form three clusters (Fig. 1B). Of the three frequently mutated residues, R320 and R470 sites grouped together in cluster 1, and G333C/S mutants grouped in cluster 2 (Fig. 1B). Though not much is known about these KEAP1 mutants, G333C and G333S are shown to be LOF mutants (Lee *et al*, 2009). R320Q and R470C mutants, on the other hand, are shown to be involved in increased interaction with NRF2 protein (Hast *et al*, 2014). There were also other KEAP1 mutants in clusters 1-3 with unknown functional significance. One such mutant, KEAP1 G480W was seen in both clusters 1 and 2. We next assessed the NRF2 activity score based on a NRF2 activity metric (Härkönen *et al*, 2023) in clusters 1-3. As a control, TCGA samples without any mutations in KEAP1 or NRF2 were used. We did not find any significant difference in the NRF2 activity score between different clusters (Fig. 1C), indicating that the differences in gene expression could not be explained by NRF2 activity status alone. We then conducted a gene set enrichment analysis (GSEA) to identify differentially expressed genes between cluster 1 and cluster 2 (Fig. S1) (Suppl. Table. S1). Cluster 1 showed significantly enriched oncogenic pathways such as mitotic cell cycle, E2F transcription, G2/M checkpoint, IFN-γ response, EMT, MYC targets and TNFα-NFκB signaling (Fig. 1D) (Suppl. Table. S2). Downregulated pathways include fatty acid metabolism, TGF-β signaling, and P53 pathway (Suppl. Table. S2). Since not much is known about the relationship between TNFα-NFκB and KEAP1 signaling, we further examined the enriched TNFα-NFκB pathway genes present in cluster 1. We found that several NFκB target genes were significantly enriched in cluster 1, when compared to G333 mutants (Fig. 1E). Upregulated genes in cluster 1 also include cell cycle and anti-apoptotic genes (Pahl, 1999; Ngo *et al*, 2020), such as *MKI67, PLK1, BRCA2, PLK4, CCNE1, CDK1, BIRC5* and *MCM2, 5 and 6* (Fig. 1E) (Suppl. Table. S1 & S2). The results suggest that KEAP1 mutations in cluster 1 might have an NRF2-independent role in regulating apoptosis in lung cancer cells.

**Fig. 1.**
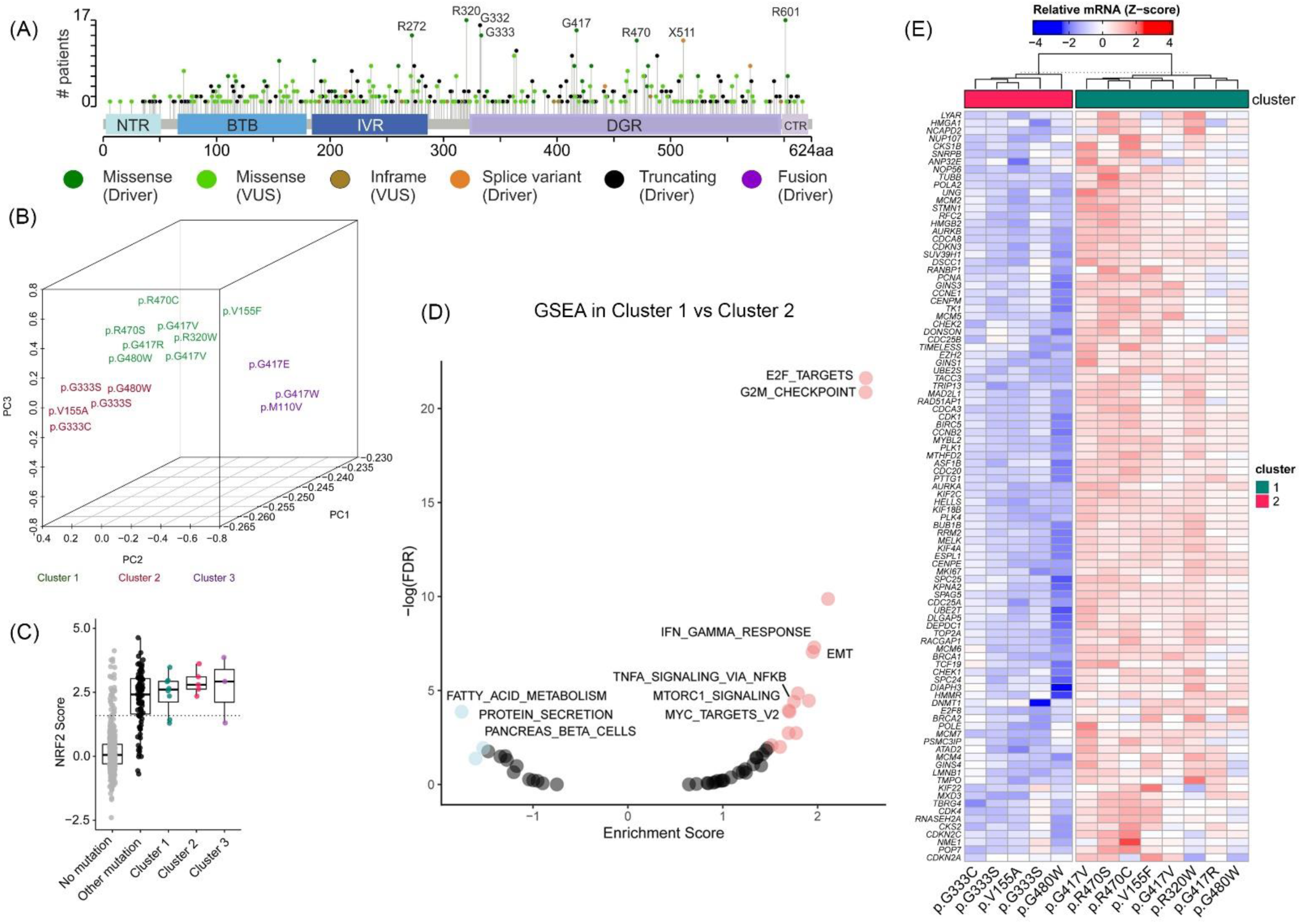
KEAP1 mutations from TCGA lung adenocarcinoma cohort classified as clusters 1-3 show differential expression of genes in oncogenic pathways. (A) Lollipop map of hotspot mutations in KEAP1 protein. (B) Clustering of KEAP1 mutations into three clusters by PCA according to their mRNA expression assessed by RNA-seq. (C) NRF2 activity score calculated in clusters 1-3. (D) Gene Set Enrichment Analysis (GSEA) from RNA-seq data between clusters 1 and 2. Positive and negative enrichment score (p < 0.05) are shown in red and blue colors, respectively. (E) Analysis of differential expression of genes between clusters 1 and 2. Data represents mean values from n = 7-9 patient samples from each KEAP1 mutant group.

### 2. Molecular modeling of KEAP1 dimers reveals that mutants R320Q and R470C show more structural flexibility to interact with other protein partners

Next, we wanted to see the molecular level differences between the native and mutant KEAP1, especially the R320Q and R470C forms. First, using molecular dynamics simulation, we characterized the interaction of native and mutant amino acid residues in R320 and R470 sites with their surrounding environment. Molecular dynamics (MD) simulation of KEAP1 R320 and R470 for one microsecond showed differences in the type of interaction with the surrounding amino acids (Fig. 2A). In native form of KEAP1, R320 residue showed a salt bridge with E242 and a hydrogen bond with main chain carbonyls of R234, A270, G242 and I232 (Fig. 2B). R470 residue showed a salt bridge interaction with D422 and a hydrogen bond with E493 from Kelch domains (Fig. 2B). In KEAP1 R320Q mutant form (shown as yellow structure in Fig. 2C), hydrogen bond interactions were shifted mainly to water bridge interactions (Fig. 2C), whereas in KEAP1 R470C mutant (shown as yellow structure in Fig. 2D), the side chain hardly formed pairwise interactions. However, in this configuration, histidine in H311 formed hydrogen bond interactions with other carboxylic acid containing residues D422 and E493 (Fig. 2D). In native form of KEAP1, MD simulations showed that H311 acts as a linker between main chain carbonyl of N489 and carboxylic acid sidechain of E493 (Fig. 2E). In KEAP1 R470C mutant form (shown as yellow structure in Fig. 2F and G), no pairwise interactions with surrounding amino acids, D422 and E493 were observed at the beginning (Fig. 2F) and end of MD simulations (Fig. 2G).

**Fig. 2.**
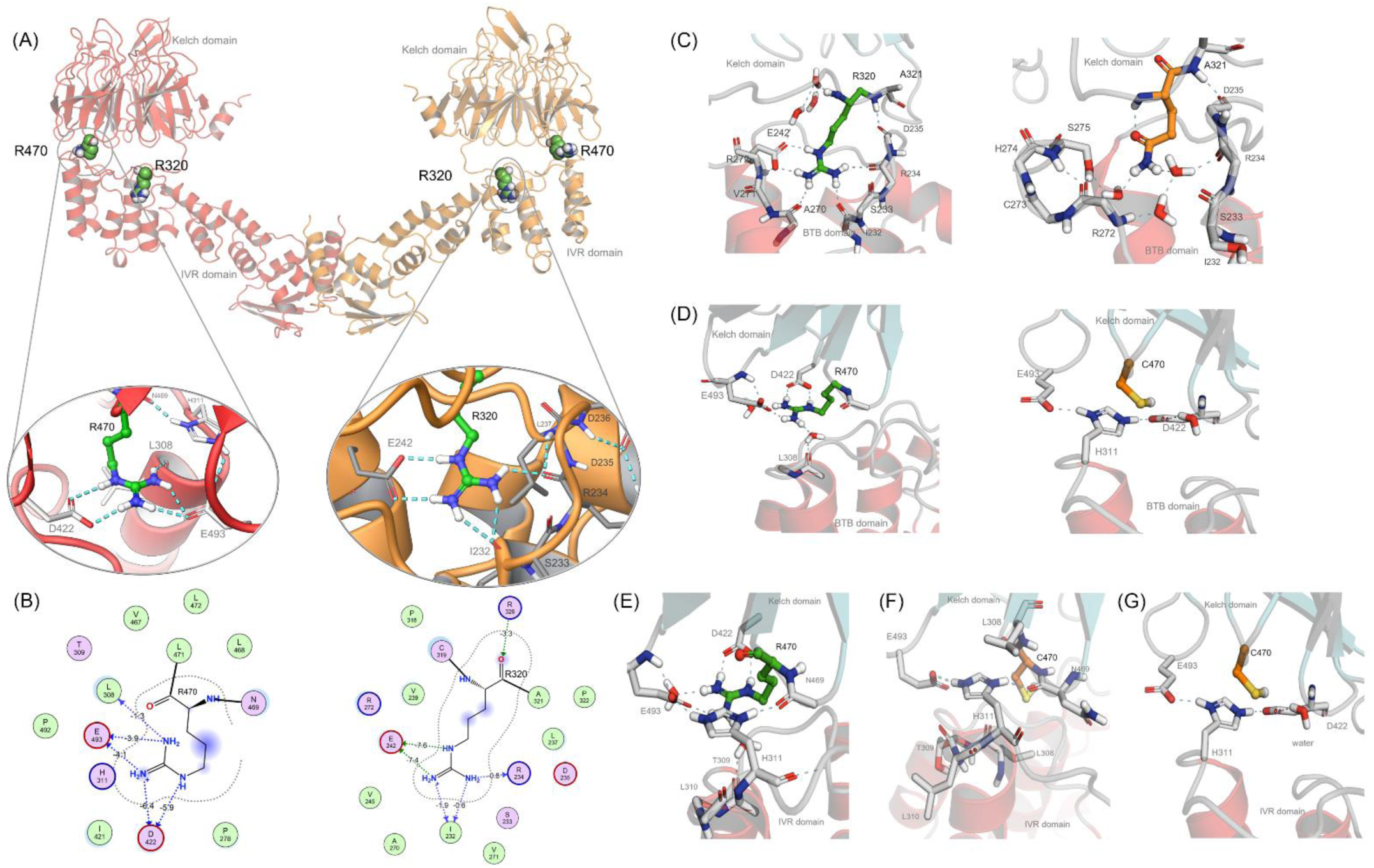
Molecular modeling and simulations of KEAP1 wt and mutants R320Q and R470C. (A) Full model of dimeric Keap1 containing BTB and Kelch domains. At the middle, pairwise interactions for native Keap1 for mutated residues R470 (left) and R320 (right) taken from AlphaFold model aftewr full protein preparation. (B) 2D interaction charts including rough estimations for hydrogen bond and ionic interaction energies according to MOE software. (C) Interactions of native (left: green) and R320Q mutant (right: yellow) of KEAP1 with surrounding amino acids. (D) Interactions of native (left: green) and R470C mutant (right: yellow) of KEAP1 with surrounding amino acids. (E) Snapshot after 1 ms of simulation shows that in native form H311 acts as linger between mainchain carbonyl (N489) and carboxylic acid sidechain of E493. (F) Snapshot shows interactions of R470C mutant at the beginning of the simulation and (G) shows interactions with D422 and E493 at the end of 1ms simulation.

Based on the information above on the interactions of amino acids from MS simulations, we next characterized dynamic motions of KEAP1 wt and its mutations, R320Q and R470C. Specifically, we calculated inter-domain interaction MMGBSA energy, using Prime module of Schrodinger suite. The results revealed that the inter-domain interaction energies were decreased 56 kcal/mol in the case of R320Q and 17 kcal/mol in the case of R470C mutations, in comparison to wt KEAP1. The weakening of energetic interaction is likely to affect dynamic motions of full dimeric KEAP1. Furthermore, we have ran 600 ns simulations for AlphaFold based full models of dimeric KEAP1 taking advantage of gpu optimized PMEMD.CUDA code and possibility to double the length of the timestep to 4 fs. Five times replicated simulations revealed that Kelch domains were structurally highly stable, while the IVR domains were highly dynamic. This instability of dimeric energy in KEAP1 mutants, R320Q and R470C could give more structural freedom and the mutants to be more flexible than the native form. However, large deviation of domain movements among replicated simulations were observed and atomic level conclusions of endpoint structures cannot be made. Increased flexibility could partly explain the stronger binding of KEAP1 R320Q and R470C with NRF2 and may have implications on other protein-protein interactions as well.

### 3. KEAP1 AP-MS proteomics show that TRAF2 is one of the significant interaction partners of KEAP1 R320Q and R470C mutants

To study the putative protein interaction partners of KEAP1 and its mutants, R320Q and R470C, we generated doxycycline inducible KEAP1 wt and mutant overexpressing Hek-TREx-293T cell lines. In the engineered cells, NRF2 protein was significantly increased in KEAP1 R320Q and R470C mutants, when compared to mock (GFP) and KEAP1 wt cells (Fig. 3A). Using affinity purification and mass spectrometry (AP-MS) analysis by combining Streptavidin-Biotin pull down method and LC-MS/MS mass spectrometry, interacting proteins were identified (Fig. 3B, C). A complete list of proteins identified and processed by SAINT algorithm is shown in Suppl. Table. S3. TNF receptor-associated factor 2 (TRAF2) was identified as one of the highest interacting proteins in KEAP1 R320Q and R470C, compared to KEAP1 wt (Fig. 3B). TRAF2 is a well-known regulator protein of TNFα-NFκB signaling, shown to be enriched in the GSEA analysis of lung cancer patients with KEAP1 mutation in sites R320 and R470 (Fig. 1D, E). TRAF2 is involved in TNFα-NFκB signaling, thus promoting cell survival. However, its signaling functions are complex and it can function as an oncogene or a tumor suppressor depending on the context (Siegmund *et al*, 2022).

**Fig. 3.**
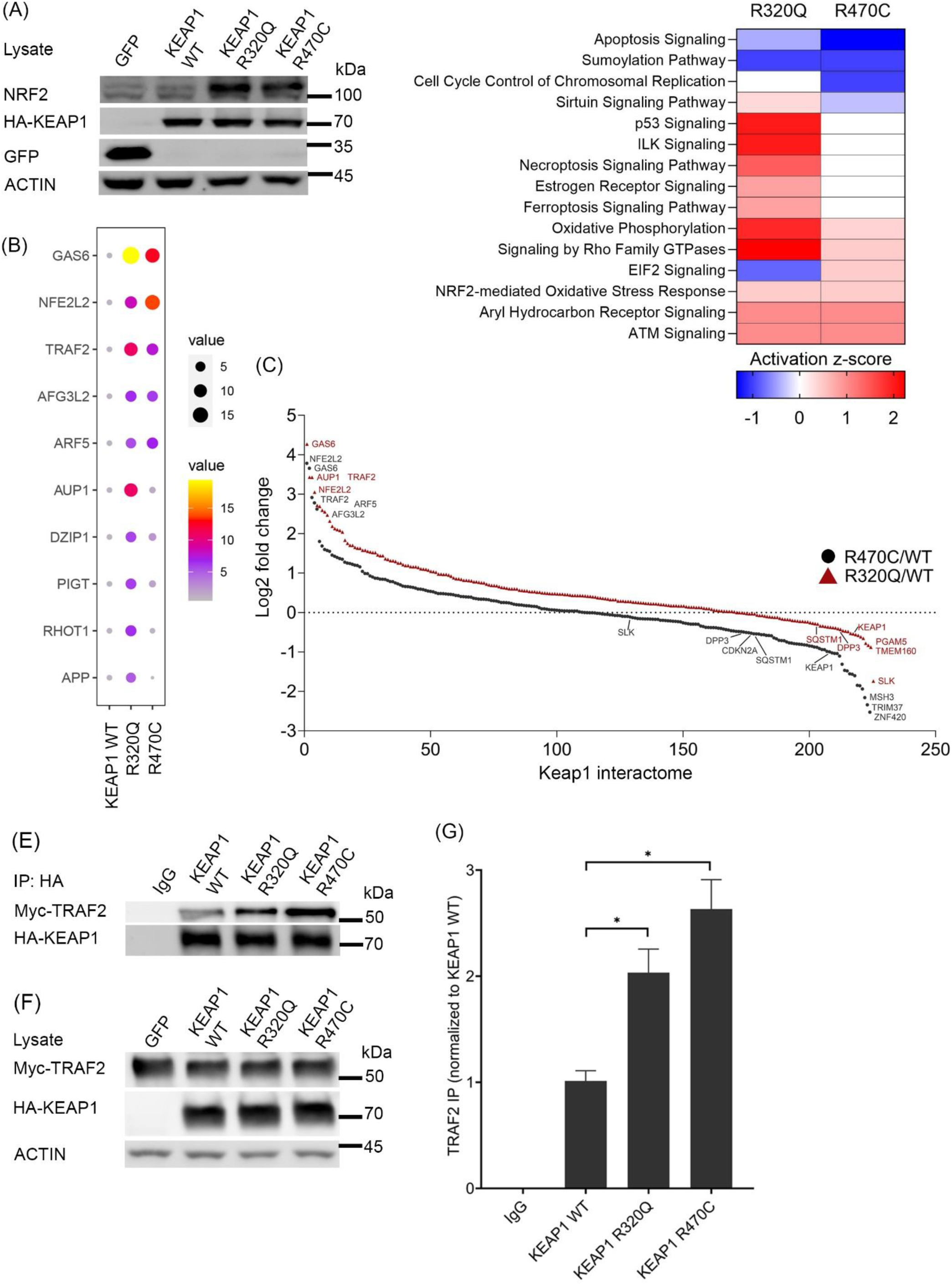
Identification of protein interaction partners of wild type KEAP1 and R320Q and R470C mutants. (A) Overexpression of HA-tagged GFP (mock), KEAP1 wild type and its R320Q and R470C mutants in Hek-TREx-293T cells. NRF2 protein expression was analyzed, with Actin as a loading control. (B) Top hits of affinity purification and mass spectrometry (AP-MS) analysis from the KEAP1 overexpressing cells. (C) Interaction partners from pathway enrichment analysis in KEAP1 mutants R320Q and R470C, as fold change to wt KEAP1. (D) Pathway enrichment analysis from AP-MS results by Ingenuity Pathway Analysis (IPA). Heatmap is based on the Z-score. (E) Co-immunoprecipation of Myc-tagged TRAF2 and HA-tagged Keap1 (F) Input controls. (G) Quantification of results from Co-IP. Data represents mean ± s.e.m from n = 3 (*p < 0.05, One way ANOVA, Tukey’s test).

Other top protein hits in the comparison between KEAP1 wt and R320Q and R470C, including NRF2 and GAS6 are listed in Fig. 3B and Suppl. Table. S3. Several well-known KEAP1 interacting proteins such as DPP3, PGAM5, SLK and P62 were identified both in KEAP1 wt and mutant proteome (Suppl. Table. S3). Most of these identified proteins, except NRF2, showed a significantly decreased interaction with KEAP1 R320Q and R470C, compared to KEAP1 wt (Fig. 3C). However, NRF2 showed a significantly increased interaction with KEAP1 R320Q and R470C (Fig. 3A, B). Apart from NRF2 and TRAF2, several anti-apoptotic proteins such as BIRC2, BIRC6, RHOT1 and 2, and MCC were also enriched in the KEAP1 R320Q and R470C protein interactome (Suppl. Table. S3). Pathway analysis of significantly enriched protein hits from AP-MS using Ingenuity Pathway Analysis (IPA) revealed that apoptosis signaling was significantly inhibited, whereas NRF2-mediated oxidative stress and oxidative phosphorylation signaling proteins were enriched (Fig. 3D). The AP-MS results of increased TRAF2 interaction with KEAP1 R320Q and R470C mutants were verified by co-IP (Fig. 3E-F). Quantification of results (Fig. 3G) showed that there is a significantly increased interaction of Myc-TRAF2 with HA-tagged KEAP1 R320Q and R470C.

### 4. Oncogenic KEAP1 mutants activate TRAF2-mediated NFκB signaling

In TNFα-NFκB signaling, TRAF2 interacts with a number of proteins such as RIPK1, BIRC2 and BIRC3, regulating apoptosis (Anderton *et al*, 2018; Vince *et al*, 2009). Upon TNFα-TNFR1 stimulus, RIPK1 acts as a scaffold for TRAF2 and TRADD complex and RIPK1 polyubiquitination leads to downstream activation of NFκB and inhibition of apoptosis (Annibaldi *et al*, 2018). However, when the TRADD-RIPK1-TRAF2 complex is disturbed, RIPK1 is deubiquitinated and its kinase activity activates caspase-mediated apoptosis pathway (Fig. 4A). BIRC2 and BIRC3 function as ubiquitin ligases to inhibit RIPK1 kinase activity and caspase activation, thereby preventing caspase-mediated apoptosis (Mifflin *et al*, 2020). Interestingly, in the GSEA analysis with the TCGA adenocarcinoma cohort, several BIRC family of genes were enriched in KEAP1 R320 and R470 mutant samples (Fig. 1E). Also, in our KEAP1 protein interactome data, the interaction of BIRC2 and BIRC6 was increased with KEAP1 mutants R320Q and R470C (Suppl. Table. S3). Additionally, there are reports indicating that RIPK1 interacts with PGAM5 which also interacts with KEAP1 (Lo & Hannink, 2006). We also identified PGAM5 as one of the significant hits interacting with wt and mutant KEAP1 (Suppl. Table. S3). We next investigated whether KEAP1 is part of RIPK1-TRAF2 complex using a pull-down assay with KEAP1 wt and mutants R320Q and R470C. Co-immunoprecipitation (Co-IP) with both Flag-RIPK1 and Myc-TRAF2 showed increased enrichment of KEAP1 mutants R320Q and R470C in comparison to wt KEAP1 (Fig. 4B and C). Quantification of KEAP1 interaction with RIPK1 and TRAF2 showed that KEAP1 mutants R320Q and R470C were enriched in both the Co-IP analysis (Fig. 4F and G).

**Fig. 4.**
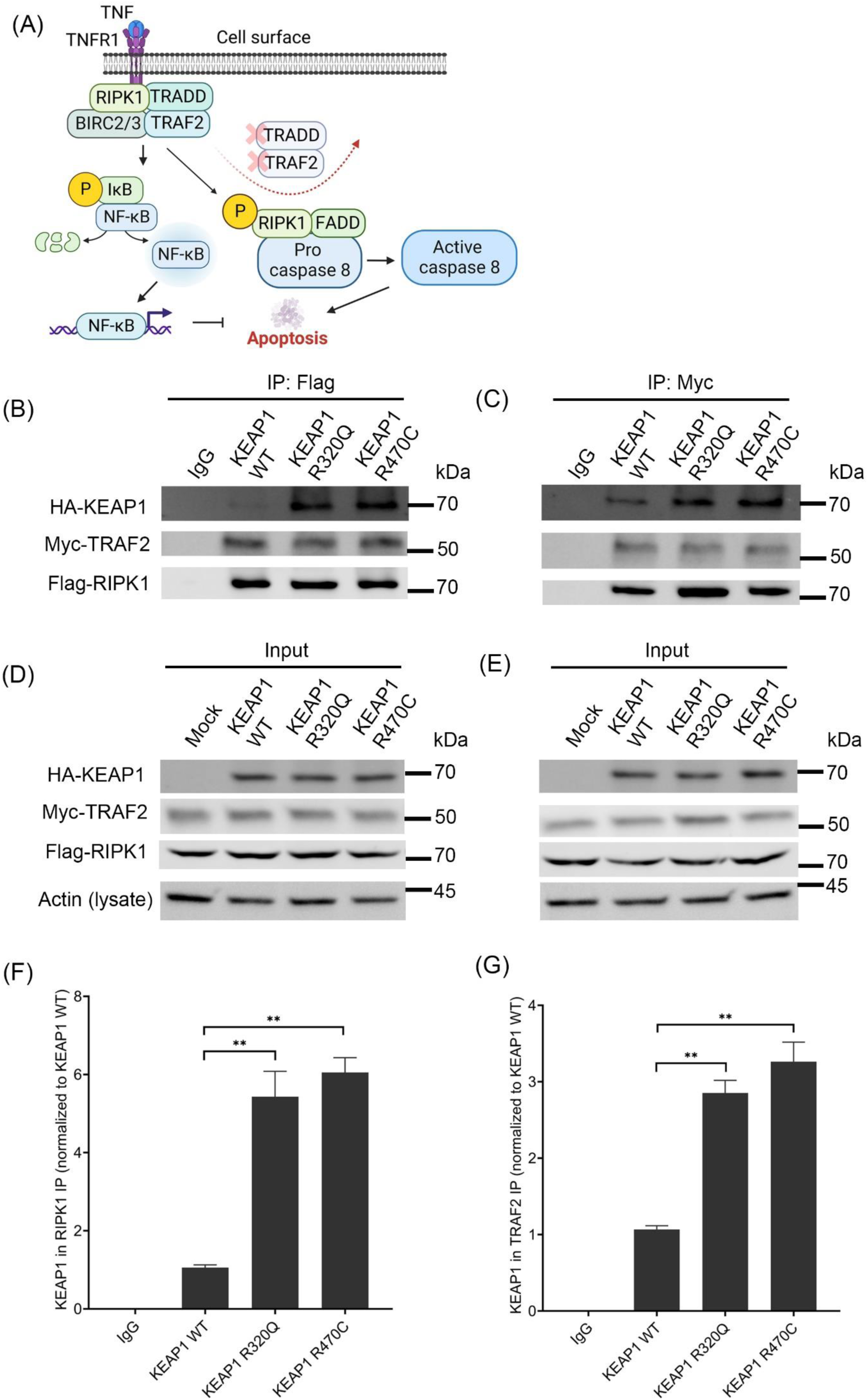
KEAP1 mutants R320Q and R470C interact with TRAF2 and RIPK1 complex. (A) Schematic illustration of TNF-TNFR1 complex proteins, consisting of TRAF2, BIRC-family of proteins, RIPK1 and TRADD. Anti-apoptotic NFκB signaling or Caspase 8-mediated apoptotic signaling is mediated by RIPK1 and its status of phosphorylation. TRAF2 is degraded and removed from RIPK1 complex to initiate apoptosis signaling. (B) A549 cells with HA-tagged KEAP1 were co-immunoprecipitated with Flag-RIPK1 or (C) Myc-TRAF2. (D) and (E) Corresponding input lysates are shown. (F) Quantification of results from Flag IP and (G) Myc IP. Data represents mean ± s.e.m from n = 3 (**p < 0.01, One way ANOVA, Tukey’s test).

Next, we investigated how KEAP1 R320Q and R470C influence the function of TRAF2. To this end, we engineered A549 cells using lentiviral overexpression (hereafter referred to as A549-LV cells) of HA-tagged mCherry (mock), KEAP1 wt, R320Q and R470C mutants (Fig. 5A). A549 cells do not have functional KEAP1, as G333C mutation found in the protein renders it inactive. KEAP1 wt transduced cells showed decreased NRF2 protein (Fig. 5A) as well as the expression of target genes NQO1, GCLM, HMOX1 and TNXRD1 (Fig. S2A-E). In contrast, KEAP1 R320Q and R470C cells showed increased NRF2 protein and target gene activation compared to mock cells (Fig. 5A) (Fig. S2A-E). Using a luciferase vector containing a minimal promoter with NFκB RELA response element, we then investigated the transcriptional activity of NFκB in A549-LV cell lines. TNFα treatment induced luciferase activity in A549-LV cell lines, with a significant increase in KEAP1 R320Q and R470C cells in comparison to mock and wild type KEAP1 controls (Fig. 5B). NFκB inhibitor, QNZ as well as two different TRAF2 specific siRNAs (Fig. S3A) inhibited the transcriptional activity of NFκB in mutant KEAP1 expressing cells (Fig. 5C). Notably, KEAP1 R320Q and R470C cells showed increased luciferase activity without TNFα stimulus, supporting the proinflammatory role of KEAP1 mutants (Fig. 5B). Inasmuch as TNFα-NFκB pathway was enriched in KEAP1 R320 and R470 adenocarcinoma samples in our GSEA analysis (Fig. 1D, E), we analyzed the expression of two NFκB target genes, IL23A and BIRC2, in transduced cells. In KEAP1 R320Q and R470C cell lines, with or without TNFα treatment, both IL23A and BIRC2 showed increased expression compared to mock and KEAP1 wt cells (Fig. S3B, C). However, with QNZ treatment, the TNFα-mediated stimulus was significantly inhibited in KEAP1 R320Q and R470C cell lines (Fig. S3B, C).

**Fig. 5.**
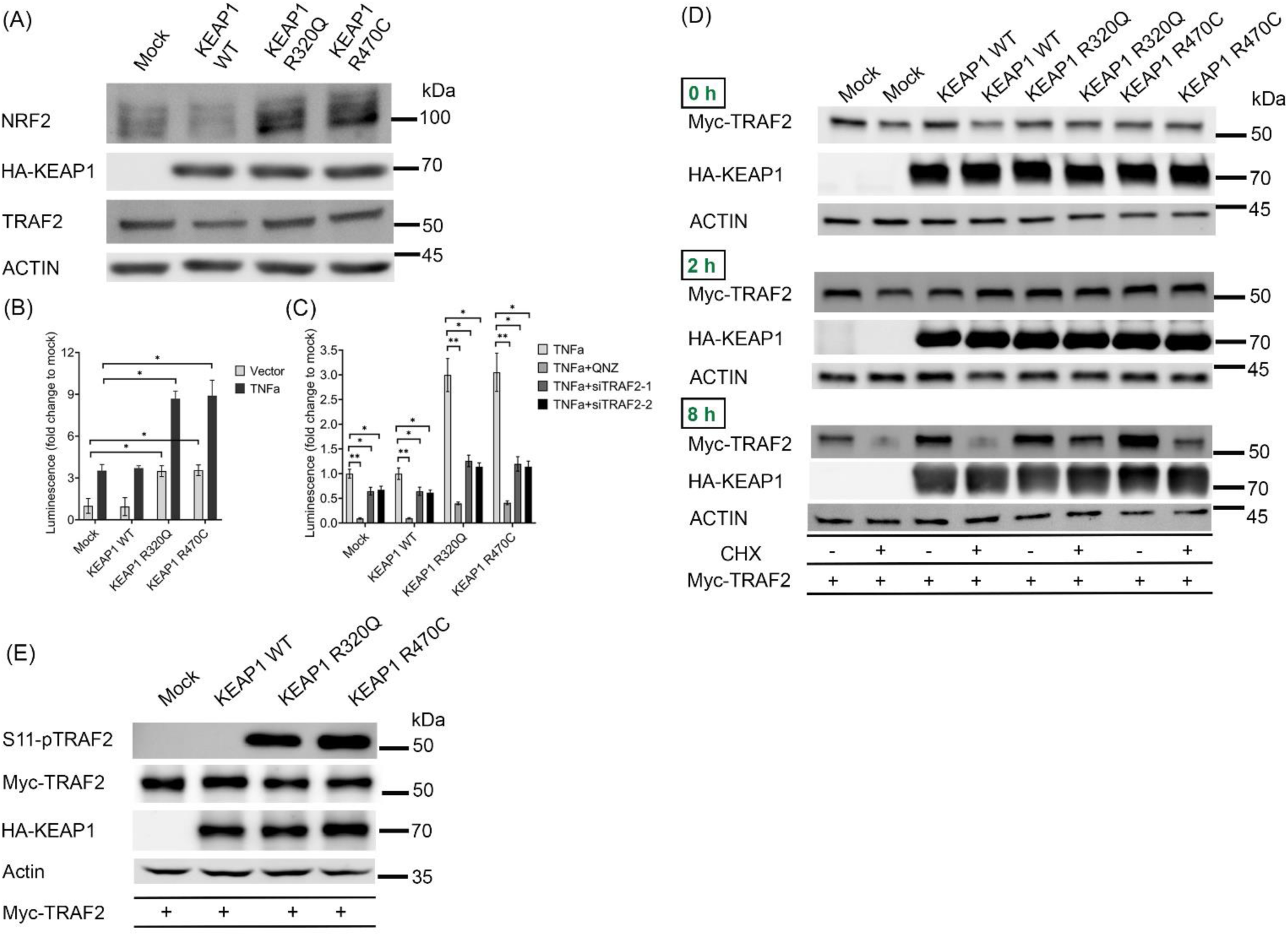
KEAP1 mutants R320Q and R470C activate NFκB transcriptional activity by stabilizing TRAF2 protein. (A) Immunoblots with respective antibodies in A549-LV cells. (B) Luciferase assay of NFκB reporter in A549-LV cells, showing NFκB transcriptional activity in KEAP1 R320Q and R470C cells with or without TNFα treatment. (C) NFκB transcriptional activity in A549-LV cells with QNZ and TRAF2 knockdown by siRNAs. (D) TRAF2 stability was studied in Hek-293T cells co-transfected with HA-KEAP1 (wt and mutants) and Myc-TRAF2 plasmids along with cycloheximide treatment for 0, 2 and 8 h. (E) Immunoblots with respective antibodies showing S11-phosphorylation of TRAF2 in A549-LV cells, transfected with Myc-TRAF2. Representative blots from three independent experiments are shown here. Data represents mean ± s.e.m from n = 3 (*p < 0.05, **p < 0.01, One way ANOVA, Tukey’s test). CHX = cycloheximide.

We then asked whether mutant KEAP1 could alter the stability of TRAF2. In the first set of experiments, Hek-293T cells co-transfected with plasmids of HA-KEAP1 (wt, R320Q and R470C) and Myc-TRAF2 were treated with protein synthesis inhibitor cycloheximide, to assess the protein stability of TRAF2 (Fig. 5D). 2 h or 8 h cycloheximide treatment was chosen on the basis of reported half-lives of TRAF2 (3-4.5 h, Habelhah *et al*, 2002) and KEAP1 (12.7 h, Taguchi *et al*, 2012). At 2 h, there was a slight decrease in Myc-TRAF2 in the mock cells, but not in other groups. After 8 h of cycloheximide treatment, there was a significant reduction of Myc-TRAF2 in the mock and HA-KEAP1 wild type transfected cells, whereas in the HA-KEAP1 R320Q and R470C cells, immunoreactivity of TRAF2 was substantially more than in the control groups (Fig. 5D). This indicates that KEAP1 R320Q and R470C could stabilize TRAF2 protein. Serine-11 (or S11) phosphorylation of TRAF2 is shown to be associated with its stability and activation (Shen *et al*, 2012). We therefore assessed S11 phosphorylation of TRAF2 in A549 cells overexpressing wt or KEAP1 mutants that were transfected with Myc-TRAF2. There was a significant increase in Myc-TRAF2 S11 phosphorylation in KEAP1 R320Q and R470C cells, but not in mock or KEAP1 wt cells. Similarly, S11 phosphorylation of TRAF2 in Hek-293T cells co-transfected with HA-KEAP1 and Myc-TRAF2 was increased in R320Q and R470C mutants (Fig. S4). The results indicate that R320Q and R470C mutants of KEAP1 increase S11 phosphorylation and stability of TRAF2, leading to activation of NFκB in lung cancer cells.

### 5. Oncogenic KEAP1 mediated TRAF2 activation leads to NFκB mediated protection against apoptosis

TRAF2 and NFκB signaling have been shown to be crucial in regulating the expression of cytoprotective genes against immune cell-mediated apoptotic signaling (Li *et al*, 2018). Inasmuch as KEAP1 R320Q and R470C mediated TRAF2 activation increased NFκB transcriptional activity, we next examined whether this prevents apoptotic cell death against chemotherapeutic agents and cytotoxic factors released by immune cells. Using confocal microscopy imaging, we first evaluated the cleaved Caspase 3/7 and propidium iodide staining in A549-LV cells, in the presence and absence of cisplatin, staurosporine and TNFα. A549 mock cells showed increased nuclear staining of cleaved Caspase 3/7 and propidium iodide on exposure to TNFα (Fig. 6A), cisplatin and staurosporine (Fig. S5A). We then measured Caspase 3/7 staining in A549-LV cells using microplate reader assay, following exposure to apoptotic agents cisplatin and staurosporine. Compared to mock and KEAP1 wt cells, there was significantly less Caspase 3/7 staining in KEAP1 R320Q and R470C cells (Fig. S5C, D). A549 mock transduced cells showed nuclear staining of cleaved Caspase 3/7 and propidium iodide when exposed to TNFα and the staining was further enhanced in cells transduced with KEAP1 wt (Fig. 6A). On the other hand, in cells transduced with R320Q and R470C mutants, no staining was observed. We also assessed PARP cleavage as an additional marker of apoptosis, with similar findings (Fig. 6B). There was less cleaved PARP in KEAP1 R320Q and R470C cells after TNFα treatment (Fig. 6B).

**Fig. 6.**
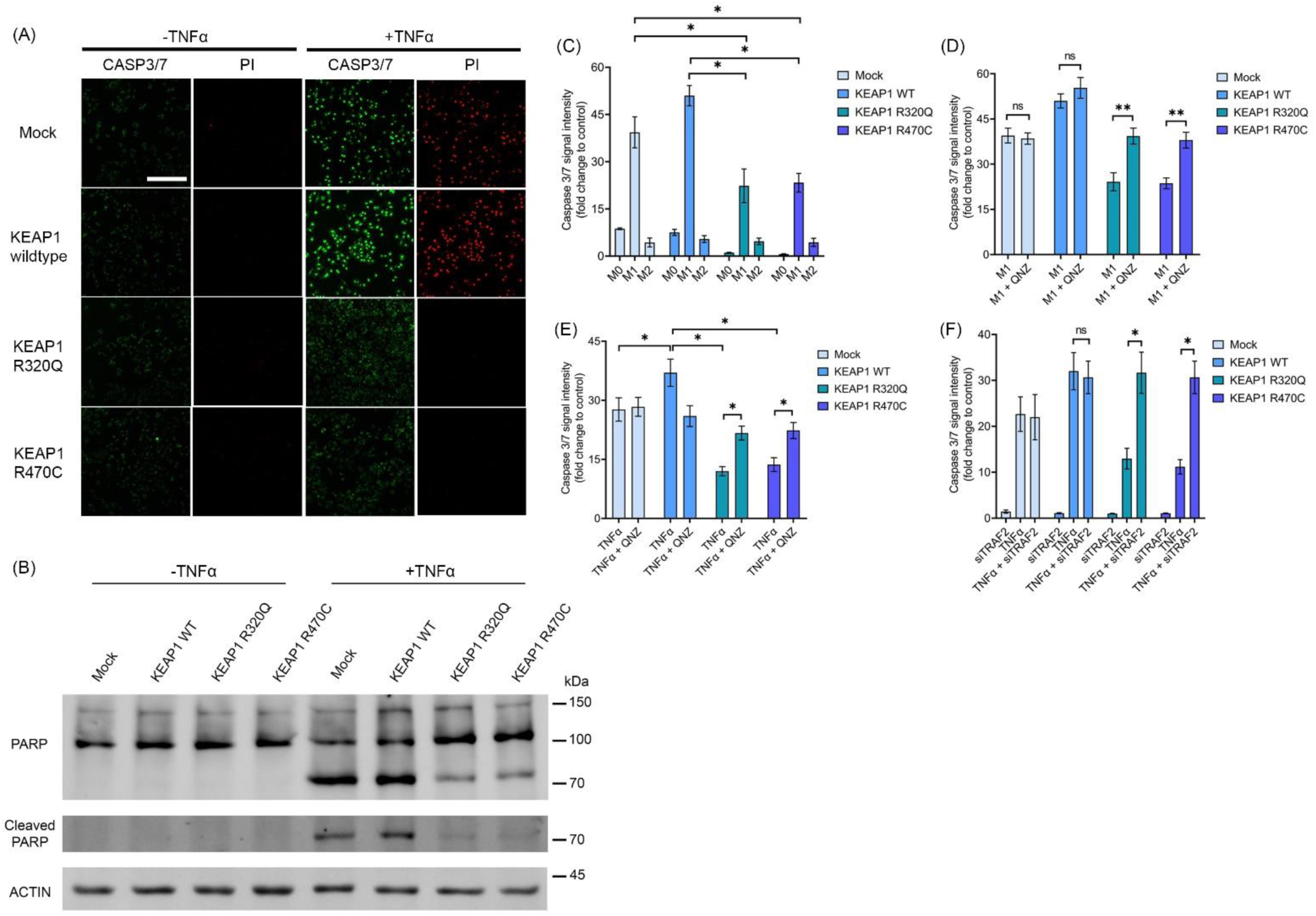
TRAF2-NFκB activation by KEAP1 R320Q and R470C leads to reduced apoptotic activity. (A) Nuclear staining of cleaved Caspase 3/7 and propidium iodide in A549-LV cells with and without TNFα treatment. Data shows representative images from n = 15 from three independent experiments. Scale bar = 100 µm. (B) Immunoblots showing full-length and cleaved PARP in A549-LV cells treated with TNFα. (C) Caspase 3/7 staining of A549-LV cells treated with THP1 CM and (D) M1 CM and QNZ. (E) Caspase 3/7 staining of A549-LV cells co-treated with TNFα and QNZ. (F) Caspase 3/7 staining of A549-LV cells with TNFα and TRAF2 knockdown. Data represents mean ± s.e.m from n = 3 (*p < 0.05, One way ANOVA, Tukey’s test).

Additionally, we assessed the effect of conditioned media (CM) of THP-1 macrophages polarized to proinflammatory M1 or immunosuppressive M2 subtypes (Kainulainen *et al*, 2022) on apoptosis in transduced A549 cells. The CM from M1 polarized THP-1 cells contains TNFα and other proinflammatory cytokines (Kainulainen *et al*, 2022). M1 CM treated cells with KEAP1 wt showed increased caspase 3/7 staining, which was substantially attenuated with KEAP1 R320Q or R470C mutants (Fig. 6C). In cells treated with M1 conditioned media (Fig. 6D) or TNFα (Fig. 6E), QNZ attenuated the anti-apoptotic effect of KEAP1 R320Q and R470C mutants. Also, TRAF2 silencing by siRNA effectively reversed the anti-apoptotic protection afforded by KEAP1 mutants R320Q and R470C (Fig. 6F). Overall, the data shows that in lung cancer cells with KEAP1 R320Q and R470C mutations, increased TRAF2 and NFκB activity is essential for the anti-apoptotic protection.

Given that KEAP1 mutants provided resistance against apoptosis caused by proinflammatory cytokines, we next examined the protein expression of 35 apoptotic markers in the presence and absence of TNFα in A549-LV cells. XIAP protein was increased in KEAP1 R320Q and R470C cells, whereas SMAC/DIABLO, TNFR1, cleaved Caspase 3 and HSP60 were downregulated (Fig. 7A, B). XIAP loss decreases the stability of TRAF2 and BIRC2 (Lawlor *et al*, 2017), and XIAP acts by inhibiting the function of pro-apoptotic RIPK1-RIPK3-Caspase-8 complex (Yabal *et al*, 2014). XIAP knockdown using siRNAs (Fig. S6) reduced NFκB RELA transcriptional activity after TNFα stimulus more robustly in KEAP1 R320Q and R470C cells (Fig. 7C). Given that Caspase 8 is upstream of Caspase 3 in the apoptotic cascade (Hughes *et al*, 2009), we next analyzed the cleaved Caspase 3/7 staining in XIAP silenced A549 transduced cells. XIAP knockdown increased TNFα-stimulated Caspase 3/7 staining in KEAP1 R320Q and R470C cells but not in mock or wt cells (Fig. 7D). Overall, the results suggest that TRAF2-NFκB activation in KEAP1 R320Q and R470C overexpressing A549 cells protects lung cancer cells against apoptosis and that XIAP and BIRC family of proteins regulate the process (Fig. 7E).

**Fig. 7.**
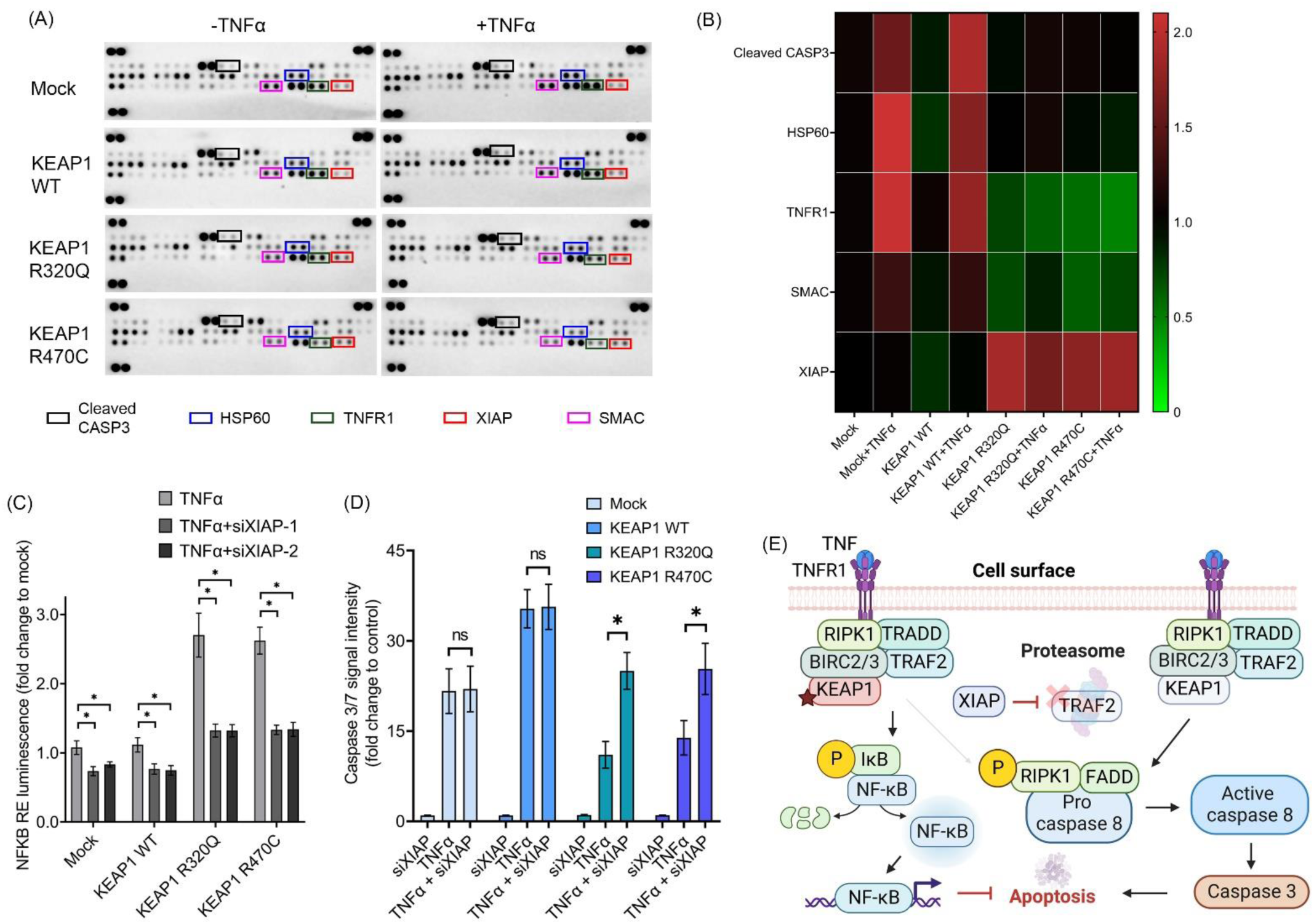
TNFα treatment in A549-LV cells show differential expression of XIAP and other key apoptotic regulators. (A) A549-LV cells treated with TNFα show differentially expressed protein markers in an apoptosis array. (B) Quantification of results from the apoptosis array. Heatmap represents average values from two dots in the array, with samples stained from three independent replicates. (C) Luciferase assay of NFκB reporter in A549-LV cells with XIAP knockdown. (D) Caspase 3/7 staining in A549-LV cells with XIAP knockdown. Data represents mean ± s.e.m from n = 3 (*p < 0.05, One way ANOVA, Tukey’s test). (E) In our hypothetical scenario, TNFα stimulated TNFR1 forms a protein complex including RIPK1, BIRC2 and 3, TRADD and TRAF2, which leads to NFκB transcriptional activity. KEAP1 mutants, R320Q and R470C (left: shown as red colored box) stabilize TRAF2 from proteasomal degradation, thereby increasing NFκB activation. Wt KEAP1 (right: shown as blue colored box) does not inhibit TRAF2 degradation thereby leading to apoptosis. XIAP, an inhibitor of TRAF2 degradation is enriched in lung cancer cells with KEAP1 R320Q and R470C mutations and contributes to anti-apoptotic signaling. Activation of TRAF2-NFκB signaling leads to increased production of cytoprotective genes, resulting in the survival of lung cancer cells against the stress induced by immune cells and chemotherapeutics. Image created with BioRender.com.

## Discussion

In NSCLC, KEAP1 and NFE2L2 mutations are highly prevalent, making the KEAP1-NRF2 signaling pathway one of the most commonly altered pathways in these cancers (Sanchez-Vega *et al*, 2018). While the GOF mutations of *NFE2L2* gene invariably occur in the KEAP1 interacting regions, KEAP1 mutations, being LOF, can occur throughout the entire sequence. Nevertheless, they are sequestered in mutation hotspots, motivating the research on their functional importance. Herein, using the TCGA data, we have identified a set of KEAP1 mutations, including hotspot mutations R320Q and R470C, that have enriched oncogenic signaling pathways (Fig. 1D). Most notably, cell cycle and anti-apoptotic pathways, such as mitotic cycle, E2F, MYC, mTORC1 and TNFα signaling pathways, were enriched. Further, we show using the affinity proteomics approach increased interaction of KEAP1 R320Q and R470C mutants with TRAF2 and other NFκB and anti-apoptotic signaling mediators, further strengthening the notion that KEAP1 signaling intersects with NFκB-mediated cytoprotective signaling. These results in conjunction with earlier observations regarding alternative substrates for KEAP1 (Ge *et al*, 2017; Taguchi *et al*, 2012) show that KEAP1 could have additional functions apart from NRF2-mediated oxidative stress regulation thereby contributing to its role in cancer.

The family of NFκB transcription factors are vital regulators of inflammation and immunity as well as cell survival and have an acknowledged role in cancer (Taniguchi & Karin, 2018; Dimitrakopoulos *et al*, 2020). NFκB activation has been shown to provide a survival advantage through induction of anti-apoptotic genes, but it may also have tumor repressor functions depending on the activation mechanism (Verzella *et al*, 2020; Taniguchi & Karin, 2018). KEAP1 has previously been proposed to be involved in suppression of NFκB activity by ubiquitination and degradation of IKK-β, which is a crucial downstream kinase regulating the NFκB pathway (Lee *et al*, 2009; Kim *et al*, 2010). In this study, we identified a novel interconnection of the NFκB and NRF2 pathways via TRAF2, a molecular adaptor that forms a protein complex with RIPK1, BIRC2 and BIRC3 upon TNFR1 receptor mediated canonical NFκB activation. TRAF2 is an oncogenic protein involved in resistance to anoikis and TNF-mediated apoptosis in cancer cells via NFκB (Vredevoogd *et al*, 2019; da Silva *et al*, 2019; Shen *et al*, 2015). Intriguingly, the loss of TRAF2 sensitizes cancer cells to checkpoint inhibition, indicating that lowering the threshold for TNF cytotoxicity augments T-cell mediated killing (Litchfield *et al*, 2021; Vredevoogd *et al*, 2019). Given that increased activity of NRF2 is associated with resistance to checkpoint inhibition (Singh *et al*, 2021), activation of the NFκB pathway via TRAF2 may provide an additional mechanism by which R320Q and R470C KEAP1 mutants could impact T-cell cytotoxicity.

TRAF2 stabilization is central to its function to support NFκB-mediated cell survival (Workman *et al*, 2020). Stabilization of TRAF2 is regulated through complex posttranslational modifications and interactions with other signaling proteins within the signalosome. TRAF2 protein has ubiquitin E3 ligase activity and is regulated through Lys63-linked autoubiquitination, necessary for the signalosome formation and NFκB activation (Zhang *et al*, 2011). Phosphorylation of TRAF2 at S11 supports K63-linked ubiquitination and NFκB activity (Workman *et al*, 2020). In this study, we show that R320Q and R470C KEAP1 mutants when coexpressed with TRAF2 increased its stability (Fig. 5D) and strongly increased the phosphorylation of S11 (Fig. 5 E), suggesting that these mutations support NFκB activation and anti-apoptotic functions of TRAF2. Given the complexity of the protein-protein interactions of TRAF2 and the regulation of its signaling, the exact mechanism by which these KEAP1 mutants interact with the TRAF2 complex to modulate its functions remains a topic of further study. Moreover, it remains to be studied whether the other KEAP1 mutants within cluster 1 (Fig 1 A) that are associated with NFκB activation have similar effects on TRAF2 signaling.

Checkpoint inhibitors have become a standard of care in NSCLC, but a large majority of patients fail to respond to the therapy. Given the protective role of TRAF2 against T-cell mediated cytotoxicity (Litchfield *et al*, 2021; Vredevoogd *et al*, 2019), inhibition of TRAF2 may provide a therapeutic option to sensitize tumors to checkpoint inhibition. However, general TRAF2 inhibition is problematic given the multifaceted role of TRAF2 in cell signaling. Therefore, therapies targeting TRAF2 interacting partners e.g., BIRC2/3 (cIAP1/2) (Vredevoogd *et al*, 2019) or selective inhibition of specific TRAF2 functions (Yan *et al*, 2022) may be more viable options. Given that KEAP1 mutations are associated with poor response to immune checkpoint inhibitors (Arbour *et al*, 2018), it can be envisioned that in KEAP1 mutant cancers with enhanced NFκB signaling such approaches could be successful.

In aggregate, we have shown that KEAP1 mutants R320Q and R470C interact with TRAF2 resulting in its stabilization and activation of NFκB in lung cancer cells. This leads to resistance to apoptosis caused by inflammatory cytokines. The study demonstrates NRF2 independent effects of KEAP1 mutations and indicates mutation-specific crosstalk between KEAP1 and NFκB signaling, highlighting the oncogenic effects of mutant KEAP1 beyond NRF2 activation.

## Materials & Methods

### Cell lines

Hek-293T and A549 cells were from ATCC, and TREx-Hek-293T cell line was obtained from Thermo Fisher Scientific. DMEM high glucose medium with glutamine (D6429, Sigma), supplemented with 10% FBS (Sigma) and 1% penicillin-streptomycin (Sigma) was used to culture the cells. Cells were regularly tested for mycoplasma contamination by PCR test (Thermo Scientific).

### Lentiviral transduction of A549 cells

Lentiviral vectors expressing mCherry-HA (mock), mCherry-HA-KEAP1 wt, mCherry-HA-KEAP1-R320Q and mCherry-HA-KEAP1-R470C were purchased from VectorBuilder. A549 cells were transduced with 10 MOI of the virus and selected for 2 weeks using 1 µg/ml puromycin in growth media. The cells expressing mCherry were then sorted with FACSAriaIII cell sorter (BD Biosciences), maintained for 2 passages and frozen until use. These transduced and sorted cells are called as A549-LV cells.

### Chemicals

TREx-Hek-293T cells with EGFP or KEAP1 overexpression were treated with 20 µg/ml cycloheximide for 2 or 8 h. A549-LV cells with mCherry or KEAP1 overexpression were treated with QNZ (0.5 µM) (Selleckchem) for 24 h, Cisplatin (25 µM) (Sigma) for 24 h, and TNF-alpha (TNFα) (10 ng/ml) (Peprotech) was used for 24 h after a pre-serum starvation (24 h). Staurosporine (0.5 µM) (Selleckchem) treatment was used for 6 h.

### Plasmids and antibodies

wt KEAP1 is cloned in N-terminally HA- and Strep-Tactin tagged pcDNA5 plasmid (Varjosalo *et al*, 2013). Site-directed mutagenesis to generate R320Q and R470C mutants from wild type (wt) KEAP1 was performed using Agilent Quick change XL mutagenesis kit. 3xMyc-TRAF2 (#44104), GFP-Ub (#11928), GFP-P65 reporter (#127172) and Flag-KEAP1 (#28023) plasmids were from Addgene. NL3.2.NF-ΚB-RE (#N1111) was from Promega. Antibodies against KEAP1, TRAF2, GFP, MYC and HA were from Proteintech. Antibody against FLAG tag was from Sigma, and antibodies against NRF2 (Proteintech) and ACTIN were from Santa Cruz. Antibodies against S11-pTRAF2, PARP and cleaved PARP were from Cell Signaling. Secondary antibodies with AlexaFluor −680, −555 and −488 labels were from Thermo Scientific.

### TCGA cohort analysis

TCGA mutation and mRNA data was downloaded from https://gdc.cancer.gov/node/905/. All data was processed with R version 4.0.0. PCA was performed with prcomp-package and k-means clustering was conducted with stats package’s kmeans function with k = 3. Differential expression analysis was conducted with edgeR and Gene Set Enrichment Analysis was performed with fgsea for the differentially expressed genes ranked by fold-change.

### KEAP1-NRF2 modeling – Protein preparation

The predicted structure containing 3D structure of Kelch-BTB domains for human Keap1 structure (Q14145) was downloaded from AlphaFold project server (Jumper *et al*, 2021; Varadi *et al*, 2022). The structure was pre-processed and minimized using the protein preparation wizard of Schrödinger Suite 2021-4 and OPLS4 force field (Protein Preparation Wizard uses modules: Epik; Impact and Prime, Schrödinger, LLC, New York, NY, 2021). Structures for point mutations R320Q and R470C were generated, and initial conformers were visually checked using rotamers library functionality of Schrodinger Maestro. Model of the dimeric human Keap1 used for graphical expressions and Amber molecular dynamics simulations was generated based on previously described AlphaFold model generation. Duplicated model was further superimposed over A and B chains of dimeric BTB domain x-ray structure (pdbid: 3I3N; (Canning *et al*, 2013)), respectively. The optimized protonation states were saved in PDB format with protonation state information.

### Molecular dynamics simulations of monomers

MD simulations were carried out using carried out using Schrödinger Desmond (Schrödinger Release 2021-4: Desmond Molecular Dynamics System, D. E. Shaw Research, Maestro-Desmond Interoperability Tools, Schrödinger, New York, NY, 2020). Prior to simulation set up the first 179 residues from BTB domain and additional helix next to Kelch domain (613-624) were removed to optimize the size of the periodic box. In Desmond system builder the orthorhombic periodic the systems were created and solvated using TIP3P waters. Systems were further neutralized including 0.1 M NaCl-salt buffer. At the beginning of the simulation, the system was subjected to default relaxation protocol of Desmond and heated up to simulation temperature of 300K. Unconstrained simulations up to length of 1 μs (microsecond) was run using NPT protocol at temperature of 300 K, pressure of 1.01325 bar, Noe-Hoover thermostat and timestep of 2 fs. Trajectories containing about 6000 snapshots were visually examined to see whether interactions between BTB and Kelch domains remain along the 1 μs. The numerical analyzes and simulation interactions charts were calculated in assistance of simulation event analysis. The prime module of Schrodinger suite was used to estimate interaction energies between separated BTB and Kelch domains. i.e. residue V316 was simply deleted to obtain separated chains. Graphical illustrations are made using either Schrodinger Maestro 2021-4 or PyMOL Molecular Graphics System, Version 2.4.1. Schrödinger, LLC. 2D-interaction plots were generated using MOE software (version 2020.09).

### Additional simulations for homodimers

Simulations were carried out using PMEMD.CUDA module of Amber18. Simulation systems were solvated using the tLeap module of Amber18 (Case *et al*, 2005) in an cuboid water box of TIP3P water molecules (Jorgensen *et al*, 1983) so that all atoms were at least 10 Å from the surface of the box. Na+ and Cl-ions were added to 0.1 M ionic concentration and amount was further adjusted to neutralize the charge of the protein. The simulation parameter files were prepared using the FF14SB force field (Maier *et al*, 2015). Finally, the “HydrogenMassRepartition” function of Parmed was used to produce the parameter files for the simulations (Hopkins *et al*, 2015). In total, three minimization steps and two steps of equilibration dynamics were performed, prior to the production simulation. The first two minimization steps, composed of 2,500 cycles of steepest descent followed by 7,500 cycles of conjugate gradient, were performed as follows: (i) in the first one, all the heavy atoms of the protein were restrained with 500 kcal/mol·Å^2^ harmonic force constant; (ii) in the second one, only the backbone atoms of the proteins were restrained 25 kcal/mol·Å^2^ force constant. The third minimization step, composed of 5,000 cycles of steepest descent and 15,000 cycles of conjugate gradient, was performed without any constraints. The equilibrium dynamics were performed in two steps: (i) 20 ps of gradual heating from 0 to 310 K under constant volume, restraining the protein atoms with 5 kcal/mol·Å2 harmonic force constant; (ii) 2 ns of unrestrained MD at constant pressure (1 bar) and constant temperature (310 K). The MD calculations employed periodic boundary conditions, the particle mesh Ewald method (Darden *et al*, 1993) for treatment of the long-range interactions beyond the 10 Å cutoff, the SHAKE algorithm (Ryckaert *et al*, 1977) to constrain bonds involving hydrogen atoms, the Langevin thermostat (Wu & Brooks, 2003) with collision frequency 1.0 ps, and a time step of 4 fs. The production MD simulations consisting of five replicates for the three systems were run for 600 ns and used the same settings as the last step of the equilibration dynamics. Trajectories (waters and ions omitted) area vailable for download at Zenodo server Gromacs binary xtc format. (https://zenodo.org/).

The equilibrium dynamics were performed in two steps: (i) 20 ps of gradual heating from 0 to 310 K under constant volume, restraining the protein atoms with 5 kcal/mol·Å2 harmonic force constant; (ii) 2 ns of unrestrained MD at constant pressure (1 bar) and constant temperature (310 K). The MD calculations employed periodic boundary conditions, the particle mesh Ewald method (Darden *et al*, 1993) for treatment of the long-range interactions beyond the 10 Å cutoff, the SHAKE algorithm (Ryckaert *et al*, 1977) to constrain bonds involving hydrogen atoms, the Langevin thermostat (Wu & Brooks, 2003) with collision frequency 1.0 ps, and a time step of 4 fs. The production MD simulations consisting of five replicates for the three systems were run for 600 ns and used the same settings as the last step of the equilibration dynamics. Trajectories (waters and ions omitted) area vailable for download at Zenodo server Gromacs binary xtc format. (https://zenodo.org/).

### Generation of KEAP1 overexpressing Hek-293T cells

TREx-Hek-293T cells were grown in 6-well plates and transfected with 5:1 ratio of pOG44 and Flp-In-FRT vectors with HA- and Strep-tagged EGFP or KEAP1 (wild type, R320Q and R470C mutants) using Fugene 6. After 48 h, cells were transferred to T175 flasks and selected using 260 µg/ml hygromycin in growth media for 2-3 weeks. Protein expression was induced using 0.5 µg/ml doxycycline (Sigma) for 24 h.

### Cell lysis

Cells grown in 10 cm plates were scraped using ice-cold PBS, spun down and treated with 0.5-1 ml of RIPA buffer supplemented with protease (Roche) and phosphatase inhibitor cocktail II (Sigma) and incubated on ice for 15 min. The samples were sonicated and then centrifuged at 14,000 x g at 4° C for 30 min. The supernatants were collected and stored in −20° C freezer until use. Protein concentration of the samples was measured using BCA kit (Pierce). 30 µg of samples were then boiled in SDS sample buffer at 95° C for 5 min and used for immunoblotting.

### Immunoblotting

30 µg of protein samples, diluted and boiled in SDS-PAGE buffer, were electrophoresed on 4-20% mini-PROTEAN TGX gels (Biorad) and transferred to nitrocellulose membrane. The membranes were blocked using 5% milk in TBST (+ 0.01% sodium azide) buffer and primary antibodies were also incubated with the same buffer. Secondary antibody staining was done in 2% milk-TBST (+ 0.01% sodium azide) and detected using ChemiDoc (Biorad).

### Co-immunoprecipitation

Cells were lysed using RIPA buffer with protease inhibitor cocktail (Roche) and in some cases with phosphatase inhibitor cocktail (Sigma) as well. 500 µg of total protein was then incubated with magnetic beads conjugated anti-HA, anti-Myc or anti-Flag antibodies according to manufacturer’s instructions (Pierce). Final elution was done using Tris-glycine elution buffer and samples were boiled using SDS-PAGE buffer at 95° C for 5 min.

### AP-MS and SAINT analysis

TREx-Hek-293T cells grown in 15cm dishes were treated with 0.5 µg/ml doxycycline for 24 h to induce protein expression and lysed with lysis buffer containing 50mM Hepes (pH 8.0), 5mM EDTA, 150mM NaCl, 50mM NaF, 0.5% IGEPAL, 1mM DTT, 1mM PMSF, 1.5mM sodium orthovanadate and protease inhibitor cocktail. Cells were kept on ice for 20 min, lysed by vortexing several times and supernatant collected after centrifugation at 16000 x g for 15 min at 4° C. Bio-Rad spin columns were prepared with IBA Strep-Tactin sepharose 50% beads and washed with lysis buffer. The lysates were then allowed to drain the column by gravity, washed three times with cold lysis buffer and biotin equilibration buffer (50mM Hepes (pH 8.0), 5mM EDTA, 150mM NaCl, 50mM NaF) and eluted with 20mM biotin (in 100mM Tris, pH 8.0) in equilibration buffer. A small aliquot of the samples was stored for immunoblotting and the rest purified using C18 columns. Trypsinized peptides were then analyzed on an EASY-nLC II system connected to an Orbitrap Elite ETD hybrid mass spectrometer (Thermo Fisher) using Thermo Scientific™ Xcalibur™ Software. A detailed description of the LC-MS/MS method used here can be found in a previous protocol (Liu *et al*, 2020). Acquired Thermo.RAW file were searched with Proteome Discoverer 1.4 (Thermo Scientific) using SEQUEST search engine of the selected human component of UniProtKB/SwissProt database. All reported data were based on peptides assigned with high confidence in Proteome Discoverer (FDR1%). The identified hits were refined by Significance Analysis of INTeractome SAINTexpress by using CRAPome and BioGRID databases, respectively, to remove contaminating proteins and to identify true hits.

### Inguinity Pathway Analysis (IPA)

SAINT Express protein hits were analyzed using Ingenuity Pathway Analysis (IPA) software (Qiagen). Significant hits were analyzed and enrichment z-score was calculated for different pathways. The interacting proteome in KEAP1 mutants −R320Q and −R470C with fold change values to KEAP1 wt was drawn using Cytoscape.

### Luciferase assay

– NFκB RE plasmid was transfected in A549-LV cells and the luciferase reporter assay was done according to Nano-Glo Luciferase Assay System protocol (Promega). TNFα was used as an activator and in some cases QNZ and TRAF2 siRNAs were also used.

### RNA silencing

Silencer select TRAF2 siRNAs (s14380, s14381) and XIAP siRNAs (s1454, s1455) were from Thermo Scientific. 30 nM of siRNA transfections were done using Lipofectamine RNAiMAX (Thermo Scientific) in Opti-MEM (Gibco), according to manufacturer’s instructions.

### qRT-PCR

RNA extraction from A549 cells grown in 6-well plates was done using High Pure RNA isolation kit (Roche). Transcriptor First strand kit (Roche) was used for cDNA synthesis and qRT-PCR was performed using FastStart Essential DNA Probes Master and gene specific probes from Universal Probe Library (Roche). Lightcycler 96 (Roche) was used for the PCR and quantification of results was done using 2^(-ΔΔCT) method. Primer sequences are given in Table 1.

**Table 1.**
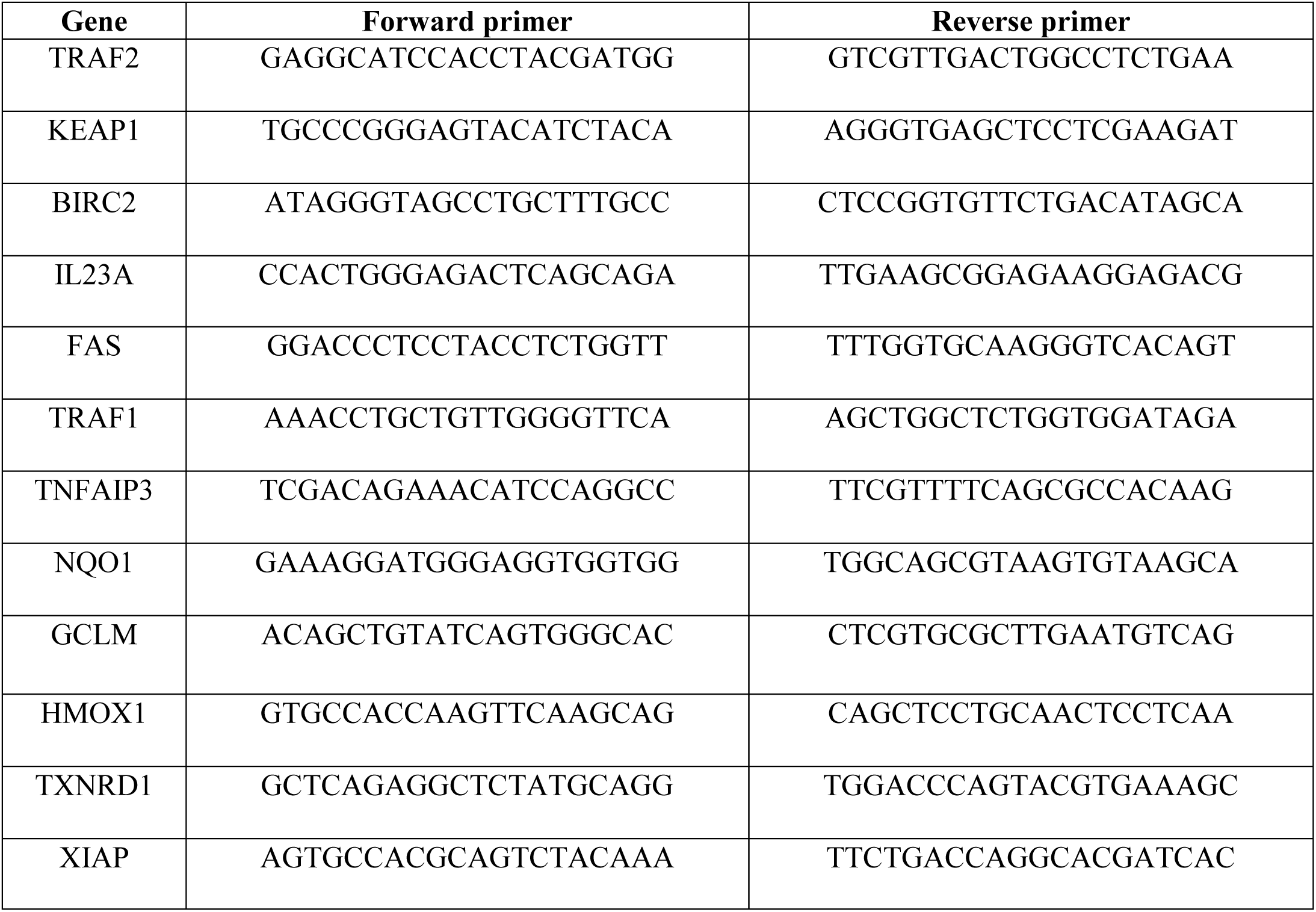
qRT-PCR primer sequences.

### Apoptosis assay

A549 cells grown in 96-well plates were stained with CellEvent Caspase 3/7 green detection reagent (Thermo Scientific) according to manufacturer’s instructions and the fluorescence measured using a plate reader at 488 nm wavelength. In some cases, full length and cleaved PARP expression were analyzed by immunoblotting.

### Live cell imaging

A549-LV cells were treated with cisplatin (25 µM), staurosporine (0.5 µM) and TNFα (10 ng/ml) for 6 h and stained for cleaved Caspase 3/7 green (5 µM) and propidium iodide (2 µM) (Thermo Scientific) for an additional 30 min. Zeiss LSM 700 confocal microscope with live cell module was used for imaging with 37° C and 5% CO_2_ conditions maintained throughout. Images were taken using a 20x objective with a numerical aperture of 0.8. The confocal images were processed using ZEN black software.

### Macrophage conditioned media (CM) treatment

THP-1 cells were differentiated into M0, M1 and M2 macrophages and conditioned media were collected as described previously (Kainulainen *et al*, 2022). A549-LV cells were treated with the CM in 1:1 ratio with growth media for 24 h.

### Proteome profiler arrays

Human apoptosis array (ARY009) and cytokine array (ARY005B) kits (R&D Systems) were used according to the manufacturer’s instructions. Briefly, 400 µg of cell lysate or 700 µl of cell culture supernatant pooled in from three independent replicates were used for each sample. The stained arrays were imaged using Bio-Rad gel documentation system and quantified using ImageJ.

### Statistical analysis

All experiments were repeated at least three times unless stated otherwise. Statistical analysis was carried out using Graphpad Prism 9.0. One way ANOVA, Tukey’s test was used with a p value < 0.05 considered significant.

## Data availability

The datasets produced in this study are available in the MassIVE databases (https://massive.ucsd.edu/) with web access MSV000092845.

## Acknowledgements

We thank Anne Karppinen, Joanna Lempiäinen and Emilia Kontio for technical assistance. Technical support from UEF cell and tissue imaging unit is also acknowledged. This study was funded by the Academy of

Finland (A-LL), Sigrid Juselius Foundation (A-LL), Finnish Cancer Society (A-LL), the Finnish Cultural Foundation (AJD), Kuopio University Foundation (AJD, SA), Horizon 2020 Framework Programme (SA, Marie Skłodowska Curie grant agreement No 740264) and Ida Montin Foundation (SA). We thank Biocenter Finland/DDCB and Biocenter Kuopio for financial support and the CSC-IT Center for Science Ltd. (Finland) for allocation of computational resources.

## Abbreviations

AP-MS: affinity purification mass spectrometry
BIRC: baculoviral IAP repeat-containing protein
BRCA2: breast cancer gene 2
CCNE1: cyclin E1
CDK1: cyclin-dependent kinase 1
CDKN1A: cyclin-dependent kinase inhibitor 1
DPP3: dipeptidyl peptidase 3
E2F: E2 factor
EMT: epithelial-mesenchymal transition
FAS: fas cell surface death receptor
GCLM: glutamate-cysteine ligase modifier subunit
GFP: green fluorescent protein
HMOX1: heme oxygenase 1
HSP60: heat shock protein 60
iASPP: inhibitor of apoptosis stimulating protein of p53
IFN-γ: interferon gamma
IKK-β: inhibitor of nuclear factor kappa-B kinase subunit beta
IL23A: interleukin-23 subunit alpha
IPA: ingenuity pathway analysis
KEAP1: kelch-like ECH-associated protein 1
LOF: loss-of-function
VUS: variants of unknown significance
MCC: MCC regulator of WNT signaling pathway
MCM: minichromosome maintenance complex component
MD: molecular dynamics
MKI67: marker of proliferation Ki-67
MTORC1: mechanistic target of rapamycin complex 1
MYC: MYC proto-oncogene
NFκB: nuclear factor kappa B
NQO1: NAD(P)H quinone dehydrogenase 1
NRF2: nuclear factor erythroid 2 - related factor 2
NSCLC: non-small cell lung cancer
P62: sequestosome-1
PALB2: partner and localizer of BRCA2
PGAM5: phosphoglycerate mutase 5
PLK: polo like kinase
RELA: RELA proto-oncogene, NF-kB subunit
RHOT1: ras homolog family member T1
RIPK: receptor-interacting protein serine/threonine protein kinase
SLK: STE20 like kinase
SMAC/DIABLO: second mitochondria-derived activator of caspase / direct IAP-binding protein with LOw pI
TGF-β: transformin growth factor beta
TNFAIP3: TNF alpha induced protein 3
TNFR1: tumor necrosis factor receptor 1
TNFα: tumor necrosis factor alpha
TRADD: tumor necrosis factor receptor 1-associated death domain protein
TRAF1: TNF receptor-associated factor 1
TRAF2: TNF receptor-associated factor 2
TXNRD1: thioredoxin reductase 1
WTX: Wilms tumor gene on the X chromosome
XIAP: X-linked inhibitor of apoptosis protein

---

**Supplementary Fig. 1.**
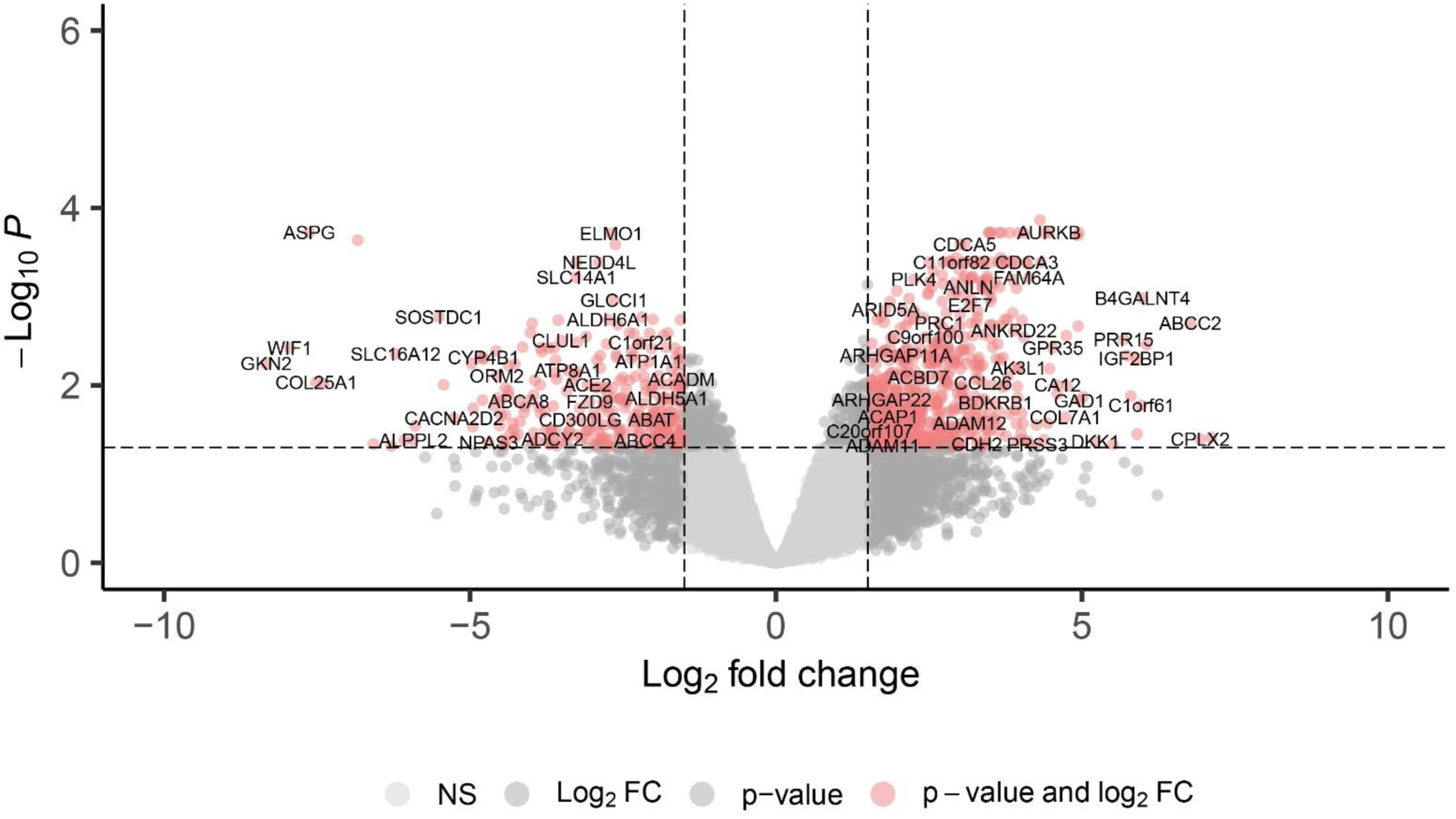
Differential expression of genes between KEAP1 mutants in cluster 1 and 2 from TCGA lung adenocarcinoma cohort. Log2 fold change and significantly upregulated and downregulated genes in cluster 1, when compared to cluster 2 KEAP1 mutant samples are shown highlighted in red color. NS = not significant. Threshold lines indicate the following: FDR-value = −log10(0.05) ∼1.3; LogFC = 1.5.

**Supplementary Fig. 2.**
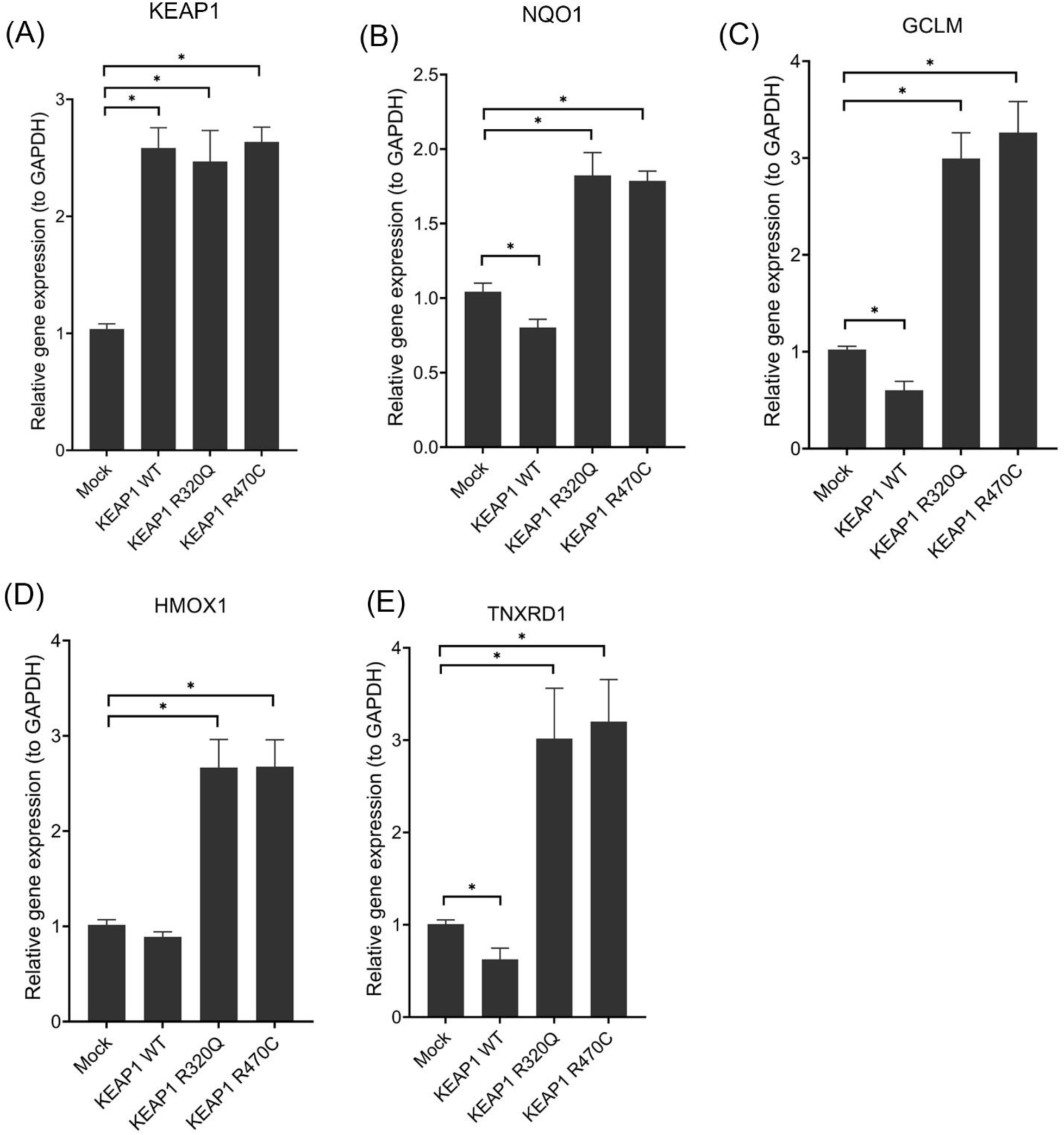
Characterization of A549-LV cells. (A) qRT-PCR analysis of KEAP1, (B) NQO1, (C) GCLM, (D) HMOX1, (E) TNXRD1 in A549-LV cells. Data represents mean ± s.e.m from n = 3 (*p < 0.05, **p < 0.01, One way ANOVA, Tukey’s test).

**Supplementary Fig. 3.**
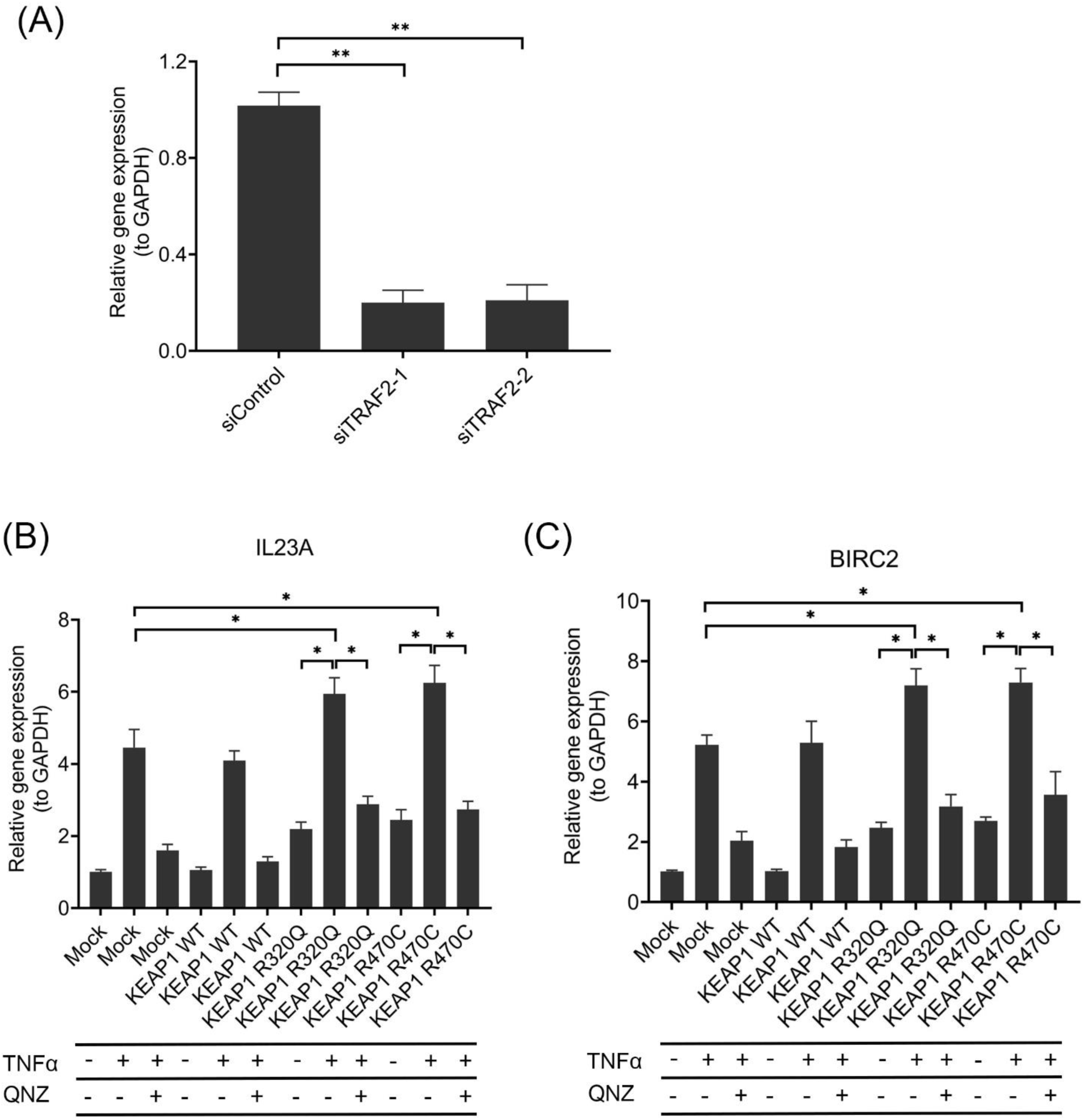
TRAF2 knockdown and qRT-PCR analysis of IL23A and BIRC2 in A549-LV cells. (A) qRT-PCR analysis of TRAF2 knockdown by siRNAs in A549 cells. (B) qRT-PCR analysis of IL23A and (C) BIRC2 genes in A549-LV cells with TNFα and QNZ treatments. Data represents mean ± s.e.m from n = 3 (*p < 0.05, **p < 0.01, One way ANOVA, Tukey’s test).

**Supplementary Fig. 4.**
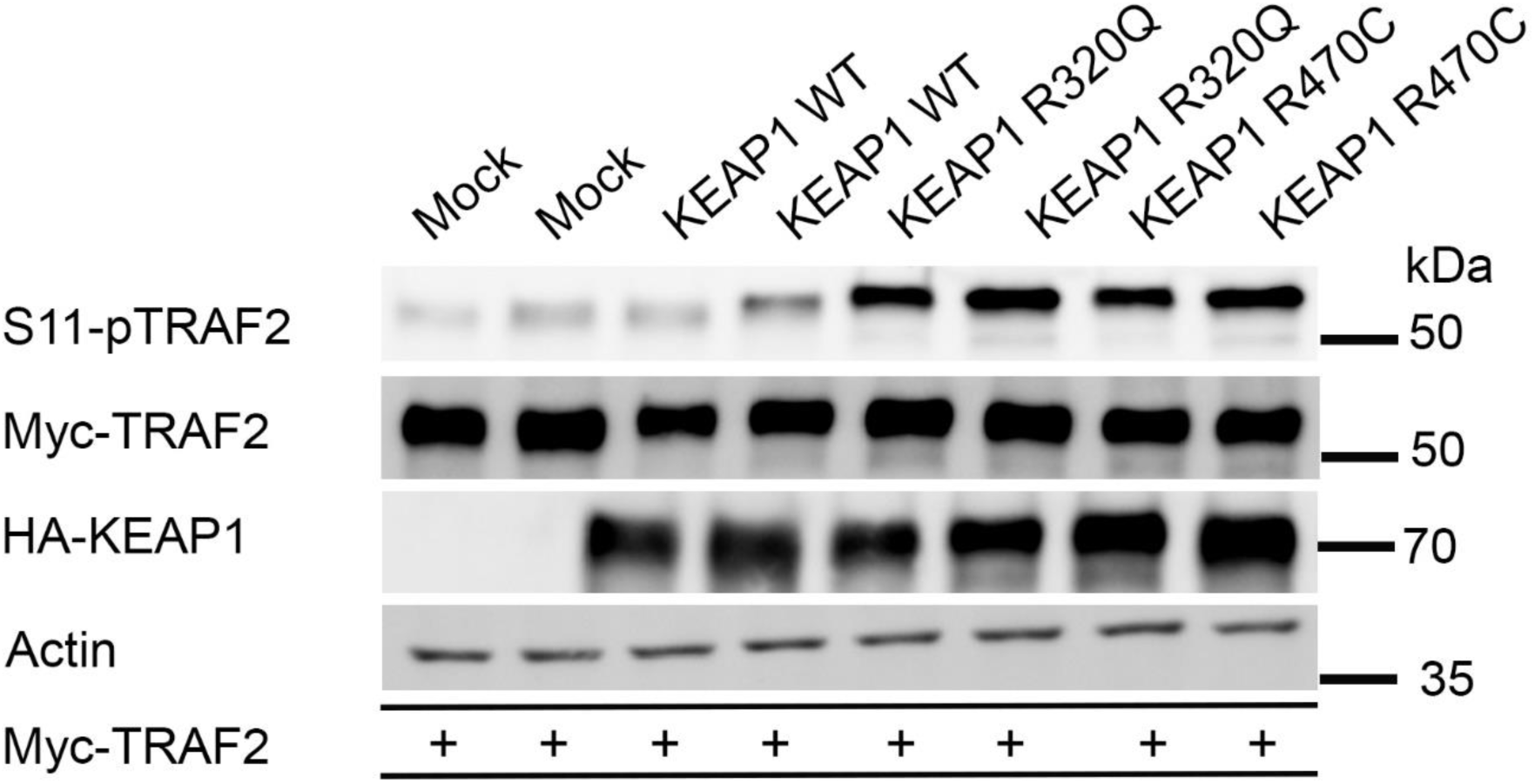
S11-phosphorylation of Myc-TRAF2 in Hek-293T cells. Hek-293T cells co-transfected with Myc-TRAF2 and HA-tagged GFP (mock), KEAP1 wt and its mutants R320Q and R470C. Immunoblots with respective antibodies are shown. Data represents blots from three independent repeats.

**Supplementary Fig. 5.**
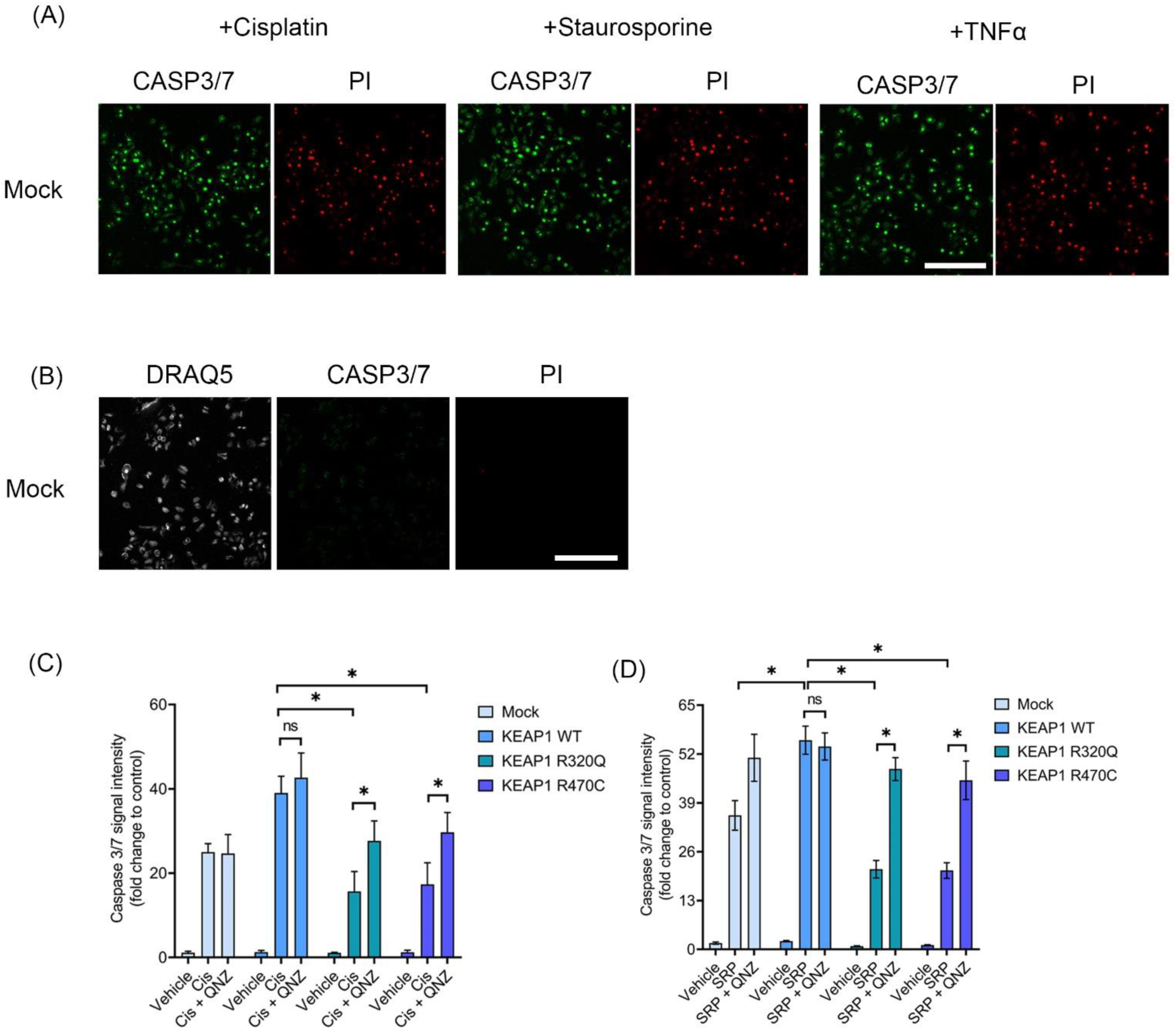
Caspase 3/7 staining of A549-LV cells. (A) A549 mock cells treated with cisplatin, staurosporine and TNFα show Caspase 3/7 and propidium iodide staining. (B) A549 mock cells without Caspase 3/7 and propidium iodide staining, showing nuclei with DRAQ5 (in white color) staining. Scale bar = 100 µm. (C) Caspase 3/7 staining of A549-LV cells with cisplatin and staurosporine, and QNZ co-treatment. Data represents mean ± s.e.m from n = 3 (*p < 0.05, **p < 0.01, One way ANOVA, Tukey’s test).

**Supplementary Fig. 6.**
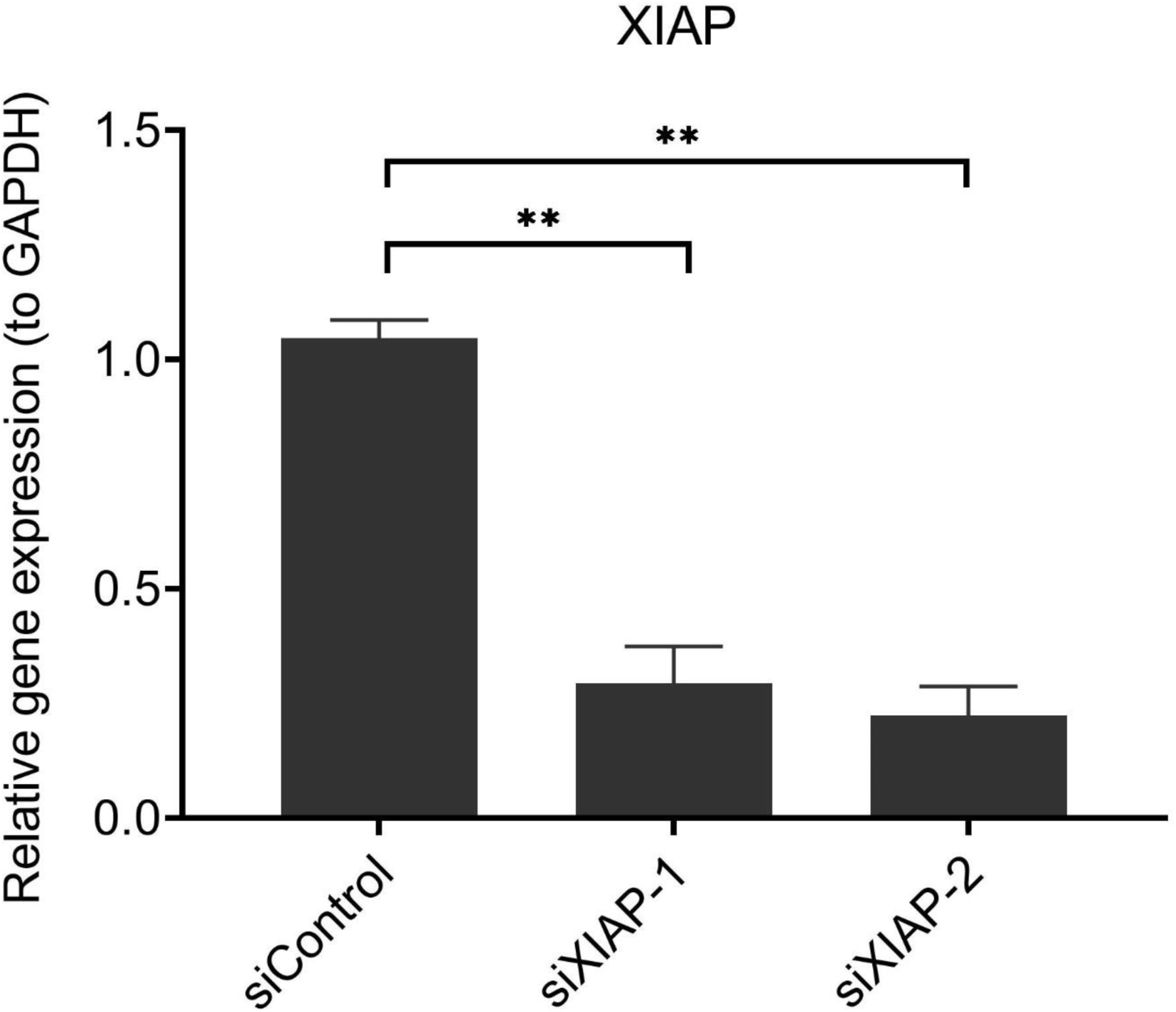
XIAP knockdown with siRNAs. qRT-PCR analysis of XIAP gene expression in A549 cells treated with siXIAP-1 and siXIAP-2. Data represents mean ± s.e.m from n = 3 **p < 0.01, One way ANOVA, Tukey’s test).

## Supplementary table legends

**Supplementary Table 1.** GSEA analysis of KEAP1 mutants in cluster 1 and 2. Pathway enrichment analysis in KEAP1 mutant samples from TCGA adenocarcinoma cohort, categorized into clusters 1 and 2.

**Supplementary Table 2** Differential expression of genes (DEGs) from GSEA analysis of KEAP1 mutants in clusters 1 and 2. DEGs in clusters 1 and 2 are shown with log2 and p values.

**Supplementary Table 3.** KEAP1 wt and mutants AP-MS and SAINT Express analysis. SAINT Express processed data on Hek-TREx-293T cells with KEAP1 wt, R320Q and R470C proteome. GFP overexpressing cells were used as mock.

## Notes

https://doi.org/doi:10.25345/C58G8FT7C

### Competing Interest Statement

The authors have declared no competing interest.

